# Effects of forced cohesin eviction and retention on X-inactivation and autosomes

**DOI:** 10.1101/2021.01.13.426565

**Authors:** Andrea J. Kriz, David Colognori, Hongjae Sunwoo, Behnam Nabet, Jeannie T. Lee

**Affiliations:** Department of Molecular Biology, Massachusetts General Hospital, Boston, Massachusetts 02114, USA; Department of Genetics, Harvard Medical School, Boston, Massachusetts 02114, USA; Department of Biological Chemistry and Molecular Pharmacology, Harvard Medical School, Boston, MA 02115, USA; Department of Cancer Biology, Dana-Farber Cancer Institute, Boston, MA 02115, USA

## Abstract

Depletion of architectural factors globally alters chromatin structure, but only modestly affects gene expression. We revisit the structure-function relationship using the inactive X chromosome (Xi) as a model. We investigate cohesin imbalances by forcing its depletion or retention using degron-tagged RAD21 (cohesin subunit) or WAPL (cohesin release factor). Interestingly, cohesin loss disrupts Xi superstructure, unveiling superloops between escapee genes, with minimal effect on gene repression. By contrast, forced cohesin retention markedly affects Xi superstructure and compromises spreading of Xist RNA-Polycomb complexes, attenuating Xi silencing. Effects are greatest at distal chromosomal ends, where looping contacts with the *Xist* locus are weakened. Surprisingly, cohesin loss created an “Xi superloop” and cohesin retention created “Xi megadomains” on the active X. Across the genome, a proper cohesin balance protects against aberrant inter-chromosomal interactions and tempers Polycomb-mediated repression. We conclude that a balance of cohesin eviction and retention regulates X-inactivation and inter-chromosomal interactions across the genome.

## INTRODUCTION

Mammalian genomes consist of meters of DNA which must be packed into a nucleus of only micrometers in diameters. Classical microscopy studies revealed that chromosomes fold into large scale structures and occupy discrete nuclear territories in order to fit into this space (Cremer and Cremer, 2001; Fraser et al., 2015; Paulson and Laemmli, 1977). The advent of genome-wide chromosome conformation capture (3C)-based assays revealed how chromosomes are organized at a finer scale. In particular, mammalian chromosomes form “loops” at several levels (Paulson and Laemmli, 1977; Rao et al., 2014; Splinter et al., 2006) and partition into “compartments” based on chromatin properties and gene expression states (Rowley and Corces, 2018). Chromosomes have also been proposed to fold into “Topologically associating domains” (TADs)(Dixon et al., 2012; Nora et al., 2012), but single cell Hi-C and microscopy studies have revealed that TADs represent population-based averages, rather than stable domains (Beagrie et al., 2017; Bintu et al., 2018; Nagano et al., 2013; Nagano et al., 2017; Stevens et al., 2017). Chromosome architecture is known to be regulated dynamically by two architectural factors — cohesin and CTCF. Cohesin is a multi-subunit complex that is thought to form a ring-like structure around chromatin and extrudes chromatin loops in an ATP-dependent manner (Davidson et al., 2019; Kim et al., 2019). RAD21 is the essential kleisin subunit of the cohesin complex, which additionally consists of SMC proteins SMC1, SMC3, and either SA1 or SA2 (Nasmyth and Haering, 2009; Peters et al., 2008). On the other hand, CTCF, an 11-zinc finger DNA-binding protein, insulates enhancers from promoters and typically anchors long-range loops (Bell and Felsenfeld, 2000; Rao et al., 2014). Importantly, interactions still cross over borders both at short range and through longer range interactions (Luppino et al., 2020).

As initial studies suggested that ectopic enhancer targeting could be induced by deletion of domain borders containing CTCF sites (Lupianez et al., 2015), it was hypothesized that these 3D genome structures had an important role in gene regulation (de Wit et al., 2015; Guo et al., 2015; Merkenschlager and Nora, 2016). Unexpectedly, though, a series of loss-of-function studies targeting architectural factors CTCF, cohesin component RAD21, and cohesin release factor WAPL showed that depletion of each of these factors dramatically alters 3D genome structure, but only modestly impacts gene expression (Haarhuis et al., 2017; Nora et al., 2017; Rao et al., 2017). Similar observations were made with CTCF binding site deletions for specific domains (Despang et al., 2019; Paliou et al., 2019; Williamson et al., 2019). These results beg the question of what precise role 3D genome organization plays during epigenetic regulation. A caveat is that these studies were conducted in a limited number of cell types and developmental timepoints.

Therefore, the possibility remains that 3D genome structure plays a more acute and global role during epigenetic regulation than is currently appreciated.

Here we revisit the role of cohesin-mediated architectural changes using a model epigenetic phenomenon, X chromosome inactivation (XCI) — the process by which female mammals silence or ‘inactivate’ one of their X chromosomes to balance dosage of X-linked genes with males. XCI occurs during differentiation of female mouse embryonic stem cells (mESCs). The process is mediated by Xist, a noncoding RNA which is transcribed from and spreads across the inactive X (Xi), triggering recruitment of repressive factors such as Polycomb Repressive Complexes 1 and 2 (PRC1/2), formation of heterochromatin, and gene silencing (Disteche, 2016; Lee, 2011; Starmer and Magnuson, 2009; Wutz, 2011). XCI provides one of the most extreme examples of changes in 3D genome structure. While the active X (Xa) is folded similarly to autosomes, the Xi shows greatly weakened TAD structure and appears to be devoid of A/B compartmentalization. The Xi is instead organized into two large (>50Mb) megadomains (Deng et al., 2015; Minajigi et al., 2015; Rao et al., 2014) and the A/B compartments are reorganized into “S1” and “S2” compartments before assuming the super-structure that characterizes the globally silenced chromosome (Wang et al., 2019; Wang et al., 2018). The restructuring from A/B to S1/S2 compartments requires Xist RNA and Polycomb repressive complex 1 (PRC1), and the further reorganization of S1/S2 into a single large compartment requires the noncanonical SMC protein, SMCHD1 (Gdula et al., 2019; Wang et al., 2019; Wang et al., 2018), a cohesin and condensin relative (Blewitt et al., 2008).

While it is now understood that SMCHD1 and the S1/S2 structures are required for gene silencing, a remaining pivotal question is whether the accompanying changes in loop domains and compartments on the Xi are required for or are merely a byproduct of silencing the entire chromosome. These larger questions impact general understanding of 3D genome regulation, as well as XCI biology. To address these questions, here we perturb cohesin levels through targeted degradation of a cohesin component RAD21 and a negative regulatory factor of cohesin, WAPL, using the dTAG system. WAPL is a cohesin release factor that evicts cohesin from DNA in prophase (Gandhi et al., 2006; Kueng et al., 2006) as well as during interphase (Tedeschi et al., 2013), possibly by opening a DNA ‘exit gate’ (Buheitel and Stemmann, 2013; Chan et al., 2012; Huis in ’t Veld et al., 2014). By degrading either RAD21 or WAPL, we modulate cohesin levels upwards and downwards in a mouse embryonic stem cell (mESCs) model and examine the consequences for loop formation, compartments, and XCI.

## RESULTS

### Toggling cohesin binding using a RAD21- and WAPL-dTAG degron system

To address how cohesin levels regulate chromosome structure and gene regulation genome-wide, and in particular during XCI, we employed the dTAG degron system (Nabet et al., 2018; Weintraub et al., 2017) to acutely and selectively degrade a cohesin component RAD21 and cohesin release factor, WAPL, in female TST ES cells (**Fig.1A**). The female TST ES line carries a *Tsix* termination cassette on one X chromosome (*Tsix^TST/+^*) to ensure silencing of the *Mus musculus* X chromosome in this hybrid ES line (Ogawa et al., 2008) (**Fig.1A**). In the dTAG system, the heterobifunctional dTAG molecule (dTAG-13) recruits an endogenous E3 ubiquitin ligase, Cereblon, to degrade FKBP12^F36V^ (herein referred to as FKBP^degron^)-tagged proteins. The dTAG degron system thus avoids previously described issues with auxin degrons (Nora et al., 2017), namely difficulties maintaining Tir1 expression and effective degradation during differentiation. The FKBP^degron^ tag was knocked in at the endogenous STOP codons of the *Rad21* and *Wapl* genes using CRISPR/Cas9 and a homology-directed repair template (**Fig.S1A-E**). We isolated 2 RAD21-FKBP^degron^ clones with comparable expression levels to endogenous RAD21 (**Fig.1B, S1B-C**). We also isolated 2 WAPL-FKBP^degron^ clones, although these appeared to have different baseline levels of WAPL expression (**Fig.1B, S1D**). We later learned that the higher WAPL expression in clone H7 was due to an amplification of the Wapl-FKBP^degron^ gene. We therefore took extra care to characterize the second clone, H2, which was karyotypically normal but had a much lower baseline WAPL level relative to wildtype cells (**Fig.S1F**). Because clone H7 had a baseline WAPL level closer to wildtype cells, we considered it important to analyze both H7 and H2. Thus, all analyses were carried out with both clones for both RAD21 and WAPL-FKBP^degron^, with results agreeing between clones.

**Fig. 1.**
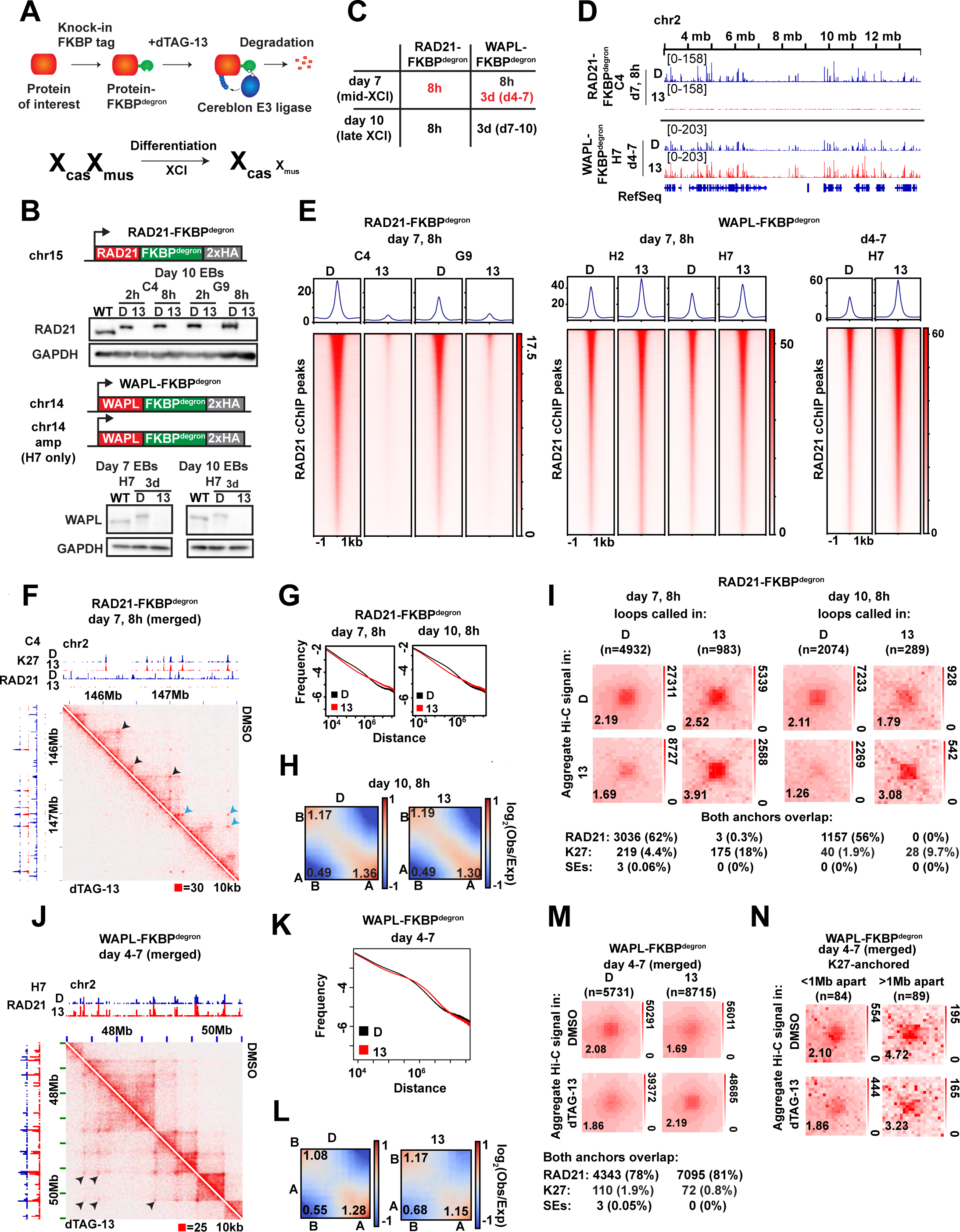
Toggling of cohesin binding and 3D genome structure during differentiation. (A) Schematic of dTAG degron system, adapted from (Weintraub et al., 2017) (top) and the TST mouse embryonic stem cell system for X chromosome inactivation (bottom). (B) Western blots for FKBP^degron^-tagged proteins and GADPH loading control in unmodified parental TST (WT) and indicated FKBP^degron^ lines. Samples treated with DMSO vehicle control (D) or 500nM dTAG-13 (13). (C) Table of time frames in each degron line investigated in this study. Red, focuses of this study. (D) Representative tracks of RAD21 binding by calibrated ChIP-seq (cChIP-seq) in RAD21-FKBP^degron^ lines and WAPL-FKBP^degron^ lines. (E) Metaplot and heatmaps of RAD21 binding by cChIP-seq in RAD21-FKBP^degron^ and WAPL-FKBP^degron^ lines. (F) Representative sections of Hi-C interaction maps in RAD21-FKBP^degron^ lines (10kb resolution, clones merged) at day 7 after 8 hours of dTAG-13 or DMSO control treatment. Cohesin-dependent (black arrows) and cohesin-independent (blue arrows) loops indicated. (G) Frequency versus distance plots of RAD21-FKBP^degron^ Hi-C maps (clones merged). Treatment as in (F). (H) Saddle plot analysis of compartment strength in RAD21-FKBP^degron^ Hi-C maps (clones merged). Scores represent B-B, B-A, and A-A compartment interaction strength. Treatment as in (F). (I) Aggregate peak analysis (APA) plots of aggregate Hi-C signal over sets of loops called in dTAG-13 or DMSO RAD21-FKBP^degron^ Hi-C maps (clones merged). Lower left: average loop enrichment (P2LL ratio). Numbers and percent of dTAG-13 and DMSO loops with both anchors overlapping RAD21, H3K27me3 cChIP peaks or mESC super-enhancers (Whyte et al. 2013) are shown. (J) Representative sections of Hi-C interaction maps in RAD21-FKBP^degron^ lines (15kb resolution, clones merged) at day 7 after 3 days of dTAG-13 or DMSO control treatment. Examples of longer cohesin-anchored chromatin loops appearing upon WAPL degradation indicated (black arrows). (K) Frequency versus distance plots of WAPL-FKBP^degron^ Hi-C maps (clones merged). Treatment as in (J). (L) Saddle plot analysis of compartment strength in WAPL-FKBP^degron^ Hi-C maps (clones merged). Scores represent B-B, B-A, and A-A compartment interaction strength. Treatment as in (J). (M) APA plots of aggregate Hi-C signal over sets of loops called in dTAG-13 or DMSO WAPL-FKBP^degron^ Hi-C maps (clones merged). Lower left: average loop enrichment (P2LL ratio). Numbers and percent of dTAG-13 and DMSO loops with both anchors overlapping RAD21, H3K27me3 cChIP peaks in corresponding samples or mESC super-enhancers (Whyte et al. 2013) are shown. (N) APA plots of aggregate WAPL-FKBP^degron^ Hi-C signal over sets of cohesin-independent H3K27me3-anchored loops (clones merged). Lower left: average loop enrichment (P2LL ratio).

Western blots and immunofluorescence confirmed that levels of RAD21 and WAPL were rapidly ablated during ES differentiation upon treatment with dTAG-13 but not with the DMSO vehicle (**Fig.1B, S1C-D, G**). To study effects on XCI, we performed analyses at days 7 and 10 of differentiation — timepoints corresponding to the mid and late stages of XCI, respectively (**Fig.1C**). For RAD21, we chose 8 hours of degradation to compare to previous degron studies (Rao et al., 2017; Rhodes et al., 2020). This degradation timeframe resulted in less impact on the cell cycle (**Fig.S2A-B**) than experienced in a previous study (Rhodes et al., 2020). For WAPL, we performed both an 8-hour and 3-day degradation, reasoning that it may take time to build up cohesin levels and observe resulting physiological effects. WAPL degradation also did not affect cell cycle profiles (**Fig.S2C-E**), consistent with the idea that separase can cleave cohesin at centromeres in the absence of WAPL, enabling cell cycle progression (Tedeschi et al., 2013).

We then performed calibrated ChIP sequencing (cChIP-seq) to examine effects on cohesin chromatin binding. RAD21 binding was reduced dramatically (**Fig.1D-E**). Whereas 29,519 RAD21 significant peaks were observed in DMSO controls, only 351 peaks were present after dTAG-13 treatment in clone C4. Similarly, whereas 23,659 peaks occurred in DMSO controls, only 780 peaks remained after dTAG-13 treatment in clone G9, Thus, there was a 99% and 97% decrease in number of RAD21 peaks in the two clones after cohesin degradation. This degree of loss in cohesin binding is comparable to a prior study (Rao et al., 2017). We also performed calibrated ChIP-seq (cChIP-seq) for RAD21 following WAPL degradation. After 8 hours of degradation, we observed increased cohesin binding (**Fig.1E**). Specifically we detected 49,130 (clone H2) and 42,086 (clone H7) peaks in DMSO control conditions and 53,480 (clone H2) and 47,866 (clone H7) peaks after dTAG-13 treatment. After 3 days of degradation, 45,415 peaks were called in DMSO control and 66,488 peaks called after dTAG-13 treatment in clone H7 (**Fig.1D-E**). In clone H2, similar numbers of peaks were called (73,706 in DMSO and 68,881 in dTAG-13), but became wider on average after dTAG-13 treatment (**Fig.S3A**). The 3-day increase in cohesin binding was similar to that observed in a previous WAPL knockout study (Haarhuis et al., 2017). Together, these data demonstrate that we can successfully toggle cohesin levels using the RAD21- and WAPL-degron system.

### Changes in 3D architecture in response to cohesin eviction and retention

To investigate changes in 3D architecture, we generated *in situ* Hi-C interaction maps, with a minimum of 360 million reads for each sample (totaling 2.8 billion reads across samples, **Table S1**) achieving a resolution of 10kb. For RAD21, 8 hours of degradation at day 7 or 10 ablated so-called topological domains evident along the diagonals of contact heatmaps (**Fig.1F-G, S3B-D**), as expected. By contrast, A/B compartmentalization was not greatly altered (**Fig.1H,S3B**), agreeing with previous studies (Rao et al., 2017; Wutz et al., 2017). On the other hand, there were intriguing effects on long-range loops. Loops are ordinarily visible in Hi-C matrices as dots of high interaction frequency (**Fig.1F, arrows**). Using HiCCUPs (Durand et al., 2016), we called 4932 loops at day 7 and 2074 loops at day 10 in RAD21-depleted cells (**Fig.1I**, DMSO columns). HiCCUPs and metaloop analysis showed a clear decrease in the number of looping interactions following cohesin depletion (**Fig.1I**) — 4,932 (DMSO) versus 983 (dTAG13) at day 7, 2074 versus 289 at day 10.

Previous studies found that ablation of cohesin dysregulates Polycomb target genes in undifferentiated mESCs (Rhodes et al., 2020), with subunits SA1 and SA2 having opposing effects on interactions between Polycomb domains (Cuadrado et al., 2019). Here, we independently observed that a substantial fraction (10-18%) of remaining loops were anchored by H3K27me3-enriched domains (**Fig.1I**). However, most cohesin-independent loops were not anchored by H3K27me3, indicating chromatin features in addition to cohesins and PRC2 are involved in loop formation (**Fig.1I, S3E**). Interestingly, metaloop analysis showed that, while the number of loops dramatically decreased, the persistent loops were strengthened (**Fig.1I**). None of the persistent loops occurred between super-enhancers (SE) (Whyte et al., 2013)(**Fig.1I**). SE interactions may occur at lower frequency during cell differentiation than in pluripotent ES cells (Rao et al., 2017; Rhodes et al., 2020). These data indicate that depleting cohesin binding reduces the number of loops across the genome, but persistent loops paradoxically have strengthened interactions.

We next examined consequences of increasing cohesin binding by degrading WAPL. When we degraded WAPL for 8 hours (to mirror the timeframe of RAD21 depletion), there was only a modest increase in cohesin binding (**Fig. 1E**). Thus, we degraded WAPL for 3 days between days 4-7 or days 7-10. These timeframes resulted in an increase in longer range interactions of 1-10Mb (**Fig.1J-K, S3B,F**) and a decrease in far-cis compartmental interactions of >10Mb (**Fig.1K-L,S3F**). Concurrently, there was a large increase in number of chromatin loops, with HiCCUPs calling 5731 loops in controls versus 8715 loops in WAPL-depleted cells, of which the vast majority were anchored by RAD21 (**Fig.1M**). Whereas RAD21-depletion strengthened Polycomb loops (**Fig. 1I**), WAPL degradation weakened them (**Fig.1N**). The effect appeared to be distance-dependent, as loops of >1Mb were weakened more than loops of <1Mb size (**Fig.1N**). Thus, whereas cohesin depletion reduces loop number and strengthens persistent loops, cohesin retention resulted in the opposite.

### Polycomb loops protect against effects of cohesin loss

We next investigated effects on gene expression by RNA-seq analysis. After 8-hr RAD21 in day 10 cells, 43 genes were significantly upregulated and 98 were downregulated (adjusted p-value < 0.1), with downregulated genes being enriched for H3K27me3 (**Fig.2A**, top, **Table S2**).

**Fig. 2.**
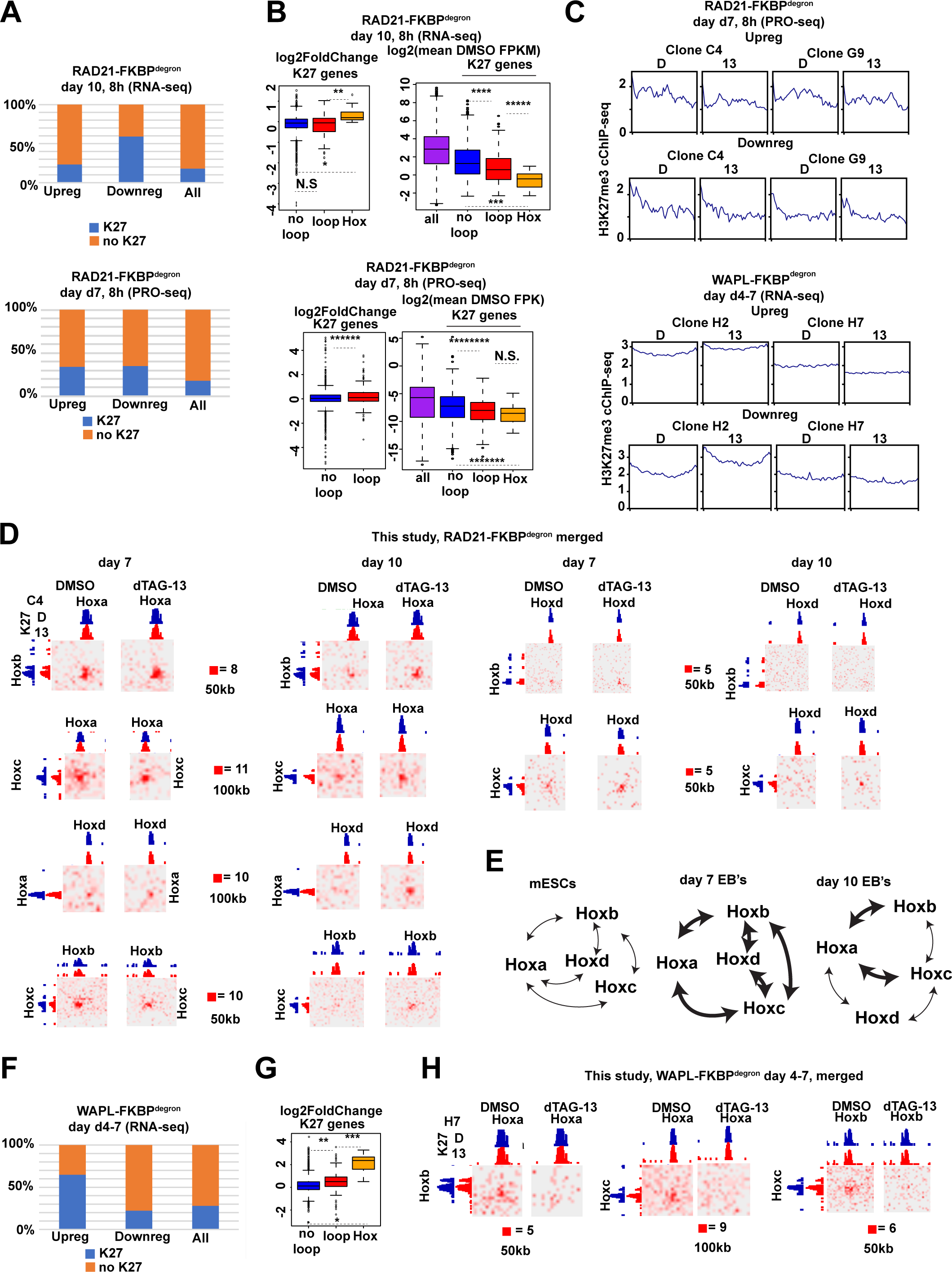
Polycomb loops protect against gene dysregulation upon cohesin loss, but increase sensitivity to cohesin gain during differentiation. (A) Genes upregulated or downregulated after indicated time courses of RAD21 degradation (relative to DMSO control) overlapping with H3K27me3 cChIP-seq peaks. All genes shown for reference. (B) RAD21-FKBP^degron^ RNA-seq (day 7: PRO-seq) DESeq2 log2-fold (dTAG-13/DMSO) change (left) or log2 FPKM (day 7: FPK, right, averaged between clones) of different categories of expressed (day 7: all) H3K27me3-marked genes. (No) loop: (not) overlapping anchor of H3K27me3-associated, cohesin-independent loop, Hox: Hox gene located in Hox cluster. All genes shown for reference. Significant p-values by Wilcoxon rank test indicated by * (p<7.344e-05), ** (p<0.0006566), *** (p<2.847e-05), **** (p<0.0005113), ***** (p<0.003082), ****** (p<0.0006716), ******* (p<0.001102), ********* (p<0.000146). N.S., not significant. (C) Average DMSO and dTAG-13 H3K27me3 cChIP-seq signal over upregulated and downregulated genes in day 7 RAD21-FKBP^degron^ and WAPL-FKBP^degron^ samples. (D) Inter-chromosomal Hi-C interaction maps between Hox cluster regions in RAD21-FKBP^degron^ lines at day 7 and day 10 at indicated resolutions (clones merged). H3K27me3 cChIP-seq signal in day 7 RAD21-FKBP^degron^ lines shown for reference (scale: 200). (E) Schematic of inter-chromosomal Hox interactions in mESCs, day 7 and day 10 of differentiation. Bold arrows indicate interactions detected by Hi-C under control conditions. (F) Genes upregulated or downregulated upon WAPL degradation from days 7-10 (relative to DMSO control) overlapping with H3K27me3 cChIP-seq peaks. All genes shown for reference. (G) WAPL-FKBP^degron^ RNA-seq log2-fold (dTAG-13/DMSO) change of different categories of expressed H3K27me3-marked genes in WAPL-FKBP^degron^ lines. Categories as in (B). Significant p-values by Wilcoxon rank test indicated by * (p<6.703e-08), ** (p<1.462e-07), *** (p<4.638e-05). (H) Inter-chromosomal Hi-C interaction maps between Hox cluster regions in WAPL-FKBP^degron^ lines (clones merged) at indicated resolutions. H3K27me3 cChIP-seq signal in day 7 WAPL-FKBP^degron^ lines shown for reference (scale: 100).

These numbers are small compared to the size of the transcriptome, but they are similar to those reported for other cell types (Rao et al., 2017; Rhodes et al., 2020). We wondered if these numbers may be an underestimate because the median mRNA half-life is ∼10 hr (Greenberg, 1972). We could have missed significant shifts in gene expression by examining steady state RNA levels. We therefore performed a transcriptome-wide analysis of nascent transcription using PRO-seq. However, a similar number of genes were revealed to be differentially expressed after 8 hr of RAD21 depletion on day 7 (**Fig.2A;** 37 downregulated, 32 upregulated), further reinforcing a modest effect of acute cohesin loss on gene expression. Loss of RAD21-loop anchors did not correlate with gene dysregulation at day 10 (**Fig.S4A**). At day 7, there was a significant tendency for genes overlapping cohesin-dependent loop anchors to be more upregulated, but again the effect was modest (**Fig.S4A**).

However, Polycomb targets inside a H3K27me3-marked persistent loop anchor were less downregulated than Polycomb targets not in such loop anchors in day 7 cells (**Fig.2B**, bottom). This contrasts with day 0 ES cells, in which Polycomb loops were correlated with increased repression upon cohesin degradation (Rhodes et al., 2020). Further examination revealed that clustered Polycomb targets tended to be more repressed than those not involved in such interactions, even in the DMSO control (**Fig.2B**). This suggests that, normally, interactions between Polycomb domains reinforce repression and cohesin depletion makes repression even more robust. Changes in gene expression upon cohesin removal were not correlated with changes in H3K27me3 by cChIP-seq (**Fig.2C**, top). Thus, Polycomb target genes are strikingly resistant to cohesin degradation in differentiating ES cells.

### Cohesin protects against inter-chromosomal *Hox* interactions and tempers Polycomb-mediated repression

Among classic Polycomb targets are genes within 4 *Hox* clusters, each located on a different chromosome, with genes in each cluster being coordinately regulated in cis to specify anterior-posterior patterning in mammals (Deschamps and Duboule, 2017; Pearson et al., 2005; Wellik, 2007). As *Hox* regulation is recapitulated during ES differentiation (Lanctot et al., 2007), we studied how architectural organization of *Hox* genes is affected by cohesin loss. Examination of Hi-C maps revealed an entire network of inter-*Hox* contacts between chromosomal clusters (**Fig.2D, S3G-I**). A small subset of these contacts had been proposed previously in ES cells (Lanctot et al., 2007; Schoenfelder et al., 2015). Heretofore unseen, however, are interactions that extend across entire *Hox* clusters and throughout cell differentiation. Most intriguingly, these trans-interactions strengthened without cohesin (**Fig.2D, S3G-I**). On the other hand, a previous RAD21-degron study did not report these contacts (Rhodes et al., 2020). Reanalysis of their Hi-C dataset indicated that they indeed exist in day 0 cells—but only after RAD21 degradation (**Fig.S3G).** Thus, analysis of our degron-tagged ES cells showed a dynamic and extensive network of *Hox-to-Hox* contacts (**Fig.2E**).

Next we asked how inappropriate cohesin retention affects these dynamics and gene expression. Because an 8-hour WAPL depletion only modestly increased cohesin retention (**Fig.S4B-D**), we depleted WAPL from days 4-7 of differentiation, where we observed significant cohesin retention (**Fig. 1E**). RNA-seq revealed differential expression of 2386 genes, of which 981 were downregulated and 1405 were upregulated (adjusted p-value < 0.1, **Table S2**). WAPL degradation at a later timepoint, days 7-10, resulted in fewer DEGs, with only 39 upregulated and 19 downregulated (**Fig.S4E**). We therefore focused further analysis on the day 4-7 timecourse. As might be expected, genes that are normally anchored by cohesin did not change expression patterns when cohesins were more strongly retained (**Fig.S4A**). Intriguingly, upregulated DEGs tend to be Polycomb targets, especially those in H3K27me3-marked persistent loop anchors (**Fig.2F,G**). This was especially evident at the *Hox* loci, where WAPL depletion and consequent cohesin retention led to a loss of trans-interaction networks and marked increase in gene expression (**Fig.2G-H, S3J**). Thus, disruption of Polycomb interaction networks, both in cis and in trans, through inappropriate cohesin retention can have catastrophic effects on gene repression.

Taken together, our data indicate that cohesins play an important role in Polycomb regulation of gene expression. Insofar as Polycomb domains have an inherent tendency to cluster (Bonev and Cavalli, 2016; Denholtz et al., 2013; Joshi et al., 2015; Kundu et al., 2017; McLaughlin et al., 2019; Rhodes et al., 2020; Schoenfelder et al., 2015; Vieux-Rochas et al., 2015), we propose that a key role of cohesins is to organize local chromatin in two respects: First, to prevent ectopic long-range cis-interactions as well as inappropriate trans-interactions; and second, to temper such interactions when they occur in the normal physiological context. Indeed, when cohesin is ablated, trans-interactions strengthen between chromosomes, as exemplified by inter-*Hox* interactions. On the other hand, when cohesins are aberrantly retained, such interactions are disrupted, with a dramatic increase in expression of the Polycomb targets.

### Cohesin regulates genes expression associated with super-enhancers in a looping-independent manner

Our RAD21/WAPL degron system enabled us to revisit an important question for which there is disagreement in the field (Rao et al., 2017; Rhodes et al., 2020). Whether super-enhancer (SE) action depends on cohesin can now be re-examined from the perspective of both cohesin loss and retention. To examine proximity of DEGs to SEs, we restricted analysis to expressed genes (FPKM > 0.5), leveraged our PRO-seq data to focus on nascent transcription, and measured genetic distances from SEs to all expressed genes, downregulated DEGs, or upregulated DEGs (Fig.3A). Significantly, downregulated DEGs in cohesin-depleted cells were closer to super-enhancers (**Fig.S5A-B**). Approximately 54% of downregulated DEGs were <150kb away, and ∼29% were <20kb away (**Fig.S5B**). Overall, loss of cohesin resulted in ∼2-fold downregulation of these targets. To ask whether this could be due to an associated shift in looping interactions, we examined Hi-C maps for interactions between SEs. Meta-loop analysis using APA plots revealed a slight enrichment of aggregate signal over SE anchors, suggesting low-frequency contacts between SEs (**Fig.S5C**). However, the contacts were not appreciably disrupted by cohesin loss in either day 7 or day 10 cells. These data contrast with those in day 0 ES cells, where SE interactions are cohesin-dependent (Rhodes et al., 2020). We conclude that cohesins play role in regulating SE-associated genes, but the effects are independent of looping in differentiating ES cells.

**Fig. 3.**
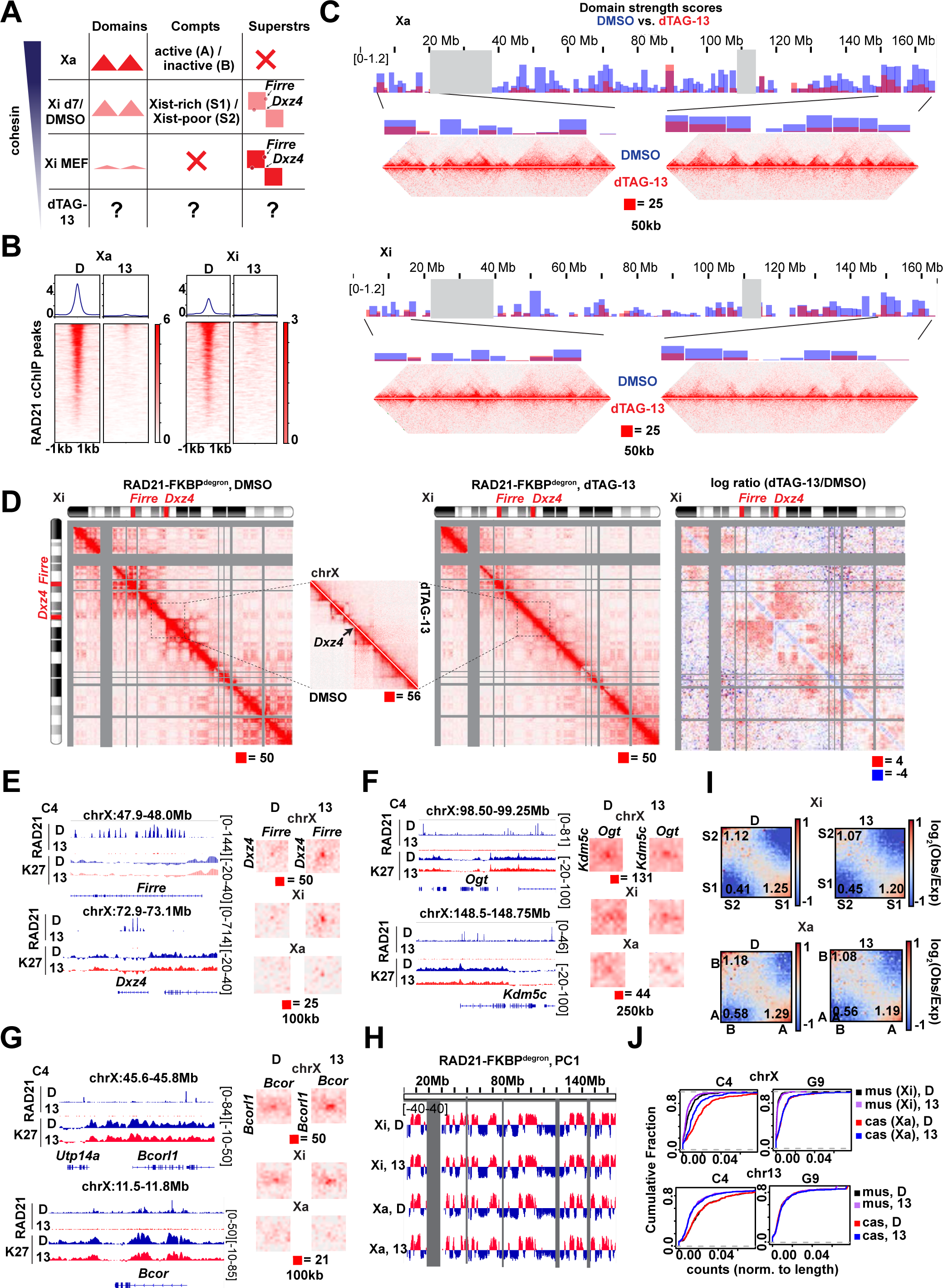
Cohesin significantly contributes to proper upregulation of proximal super-enhancer targets without affecting long-range super-enhancer interactions. (A) Distance to nearest super-enhancer (Whyte et al. 2013) for expressed genes upregulated or downregulated upon RAD21 degradation (relative to DMSO). Expressed genes shown for reference. Significant p-values by Wilcoxon rank test indicated by * (p<0.006057), ** (p<0.01569). N.S., not significant. (B) Day 7 RAD21-FKBP^degron^ PRO-seq and RAD21 cChIP-seq signal over representative super-enhancer targets downregulated upon 8 hours RAD21 degradation (relative to DMSO). (C) Aggregate Peak Analysis plots of aggregate RAD21-FKBP^degron^ Hi-C signal (clones merged) over regions anchored by super-enhancers. Super-enhancers segregated by distance into 5 quantiles.

### Xi superloops form between escapees in a cohesin- and Polycomb-independent manner

The inactive X chromosome (Xi) is a strong Polycomb target, becoming hugely enriched with H3K27me3 in female ES cells undergoing differentiation and XCI (Plath et al., 2003; Silva et al., 2003; Wang et al., 2001; Zhao et al., 2008). XCI normally brings about major architectural changes to the X chromosome, as Xist RNA evicts up to 90% of cohesins while it spreads along the Xi (Minajigi et al., 2015) (**Fig.3A and 4A,** Xi MEF row). Here we followed the Xi from mid (day 7) to late (day 10) XCI and indeed observed 40-50% loss by days 7 to 10 (**Fig.3B and 4B**). Changes in cohesin binding during differentiation are known to correlate with (i) suppression of “TADs” across the Xi (Wang et al., 2018), (ii) appearance of two megadomains with a strong border at the macrosatellite repeat, *Dxz4,* (Deng et al., 2015; Froberg et al., 2018; Giorgetti et al., 2016; Minajigi et al., 2015; Rao et al., 2014) and (iii) establishment of a ‘superloop’ spanning ∼30 Mb between *Dxz4* and another repeat element, *Firre* (Darrow et al., 2016; Hacisuleyman et al., 2014; Horakova et al., 2012; Rao et al., 2014) (**Fig.3A**). The superloops are not present on the Xa. Current models postulate that superloops form through loop extrusion of cohesin (Bansal et al., 2019; Bonora et al., 2018), but currently unknown is whether cohesin depletion is required to form the Xi superstructure and deposit the H3K27me3 Polycomb mark. Does cohesin binding antagonize the establishment of the Xi? Is cohesin eviction required for XCI? Does the remaining cohesin binding have a role in folding the Xi?

Our ability to toggle cohesin levels using the RAD21/WAPL degron system provided an opportunity to address these questions (**Figs. 3-4)**. Our in situ Hi-C was performed in a hybrid female ES line (*Tsix^TST/+^*) carrying a *Mus musculus* (mus) and a *Mus castaneus* (cas) X chromosome, with the mus X chromosome bearing a *Tsix* mutation that ensures inactivation of the X-chromosome in cis (Ogawa et al., 2008). We were therefore able to perform an allele-specific analysis to examine the Xa and Xi separately. In day 7 control cells, whereas the Xa showed characteristic topological domains, the Xi showed weakened (but not ablated) local domains (**Fig.3C, 4C**) and appearance of megadomains (**Fig.3D, 4D**), as expected. When cohesin was depleted by dTAG-13 treatment, the topological domains were nearly completely abolished on both Xa and Xi (**Fig.3C, 4C**). The loss of local interactions was also observed on the Xa, similar to autosomes (**Fig.4E,1F**). On the Xi, the megadomains and *Dxz4* border also disappeared in the absence of cohesin (**Fig.3D, 4D**). Thus, although cohesin becomes depleted on the Xi during the normal course of XCI, the remaining cohesin is required for folding into megadomains. Notably, the *Dxz4* domain contains a number of cohesin-binding sites (**Fig.3E**, RAD21 track) that normally persist on the Xi despite Xist-mediated cohesin repulsion (Bonora et al., 2018; Froberg et al., 2018; Giorgetti et al., 2016).

**Fig. 4.**
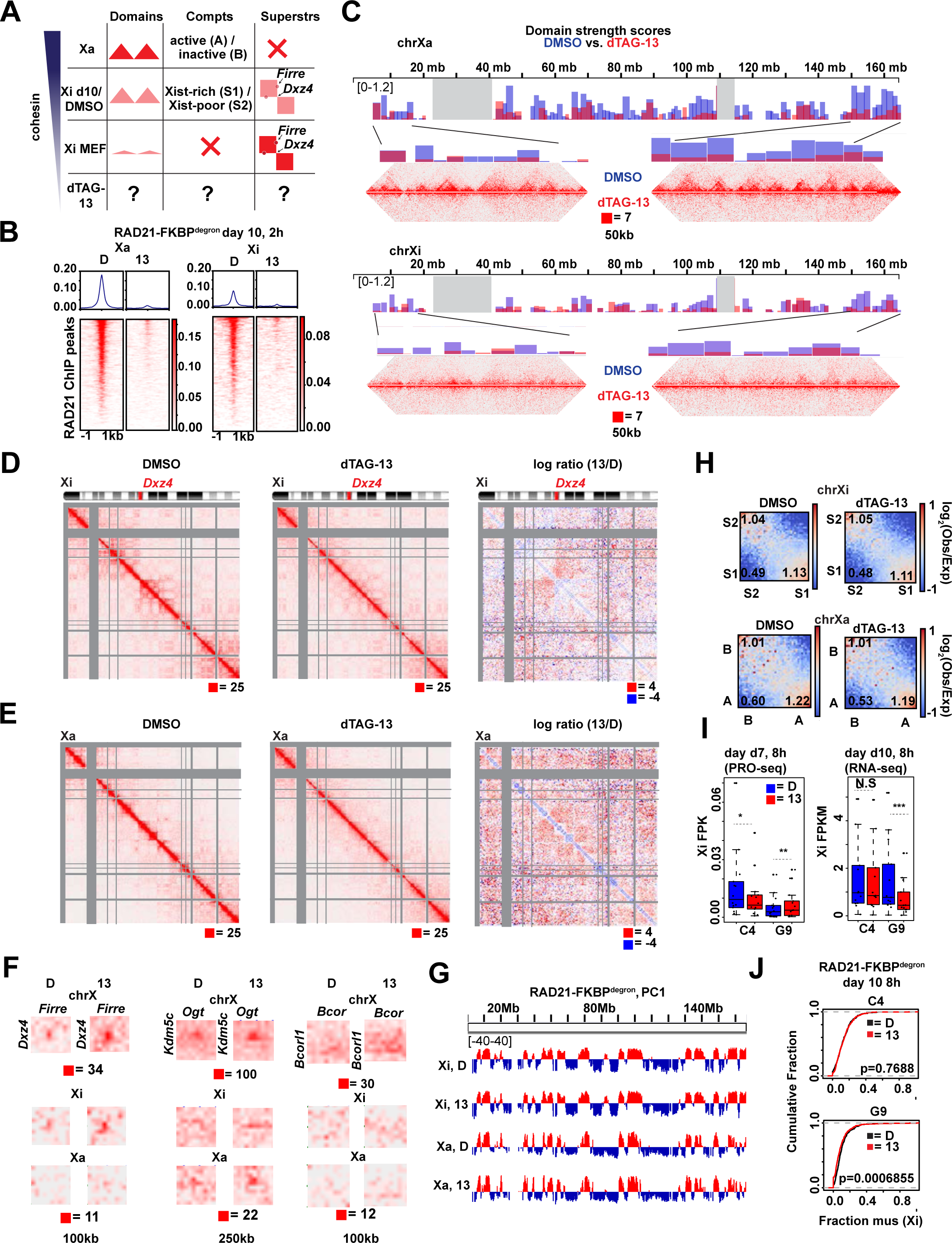
Cohesin is required for proper folding of the inactive X chromosome. (A) Schematic of expected cohesin levels versus TAD, compartment, and superstructures detected on the X chromosome at different stages of XCI, based on prior studies. Levels corresponding to day 7 DMSO control and dTAG-13 treated cells indicated. (B) Metaplot and heatmaps of active (Xa) and inactive (Xi) X chromosome RAD21 binding by cChIP-seq in day 7 RAD21-FKBP^degron^ embryoid bodies treated for 8 hours with 500nM dTAG-13 or DMSO. (C) TAD scores calculated from day 7 Xa and Xi RAD21-FKBP^degron^ allelic Hi-C interaction maps binned at 100kb. DMSO and dTAG-13 samples as in (B). Representative zoomed-in Hi-C maps shown for the indicated regions (50kb resolution, clones merged). (D) Chromosome-wide day 7 Xi RAD21-FKBP^degron^ allelic Hi-C maps (250kb resolution, clones merged) and log2-ratio map (1Mb resolution). DMSO and dTAG-13 treatment as in (B). Inset (50kb resolution) showing cohesin-dependent ‘stripe’ at the megadomain border, *Dxz4*. (E) Zoomed-in day 7 composite X, Xi, and Xa Hi-C maps at *Dxz4-Firre* superloop (100kb resolution, clones merged). RAD21 and H3K27me3 cChIP-seq at superloop anchors shown for reference. DMSO and dTAG-13 treatment as in (B). (F) Zoomed-in day 7 composite X, Xi, and Xa Hi-C maps at *Kdm5c-Ogt* loop (250kb resolution, clones merged). RAD21 and H3K27me3 cChIP-seq at loop anchors shown for reference. DMSO and dTAG-13 treatment as in (B). (G) Zoomed-in day 7 composite X, Xi, and Xa Hi-C maps at *Bcor-Bcorl1* loop (100kb resolution, clones merged). RAD21 and H3K27me3 cChIP-seq at loop anchors shown for reference. DMSO and dTAG-13 treatment as in (B). (H) First Principal Component (PC1) tracks of day 7 Xa and Xi RAD21-FKBP^degron^ Hi-C maps. DMSO and dTAG-13 treatment as in (B). (I) Saddle plot analysis of compartment strength in RAD21-FKBP^degron^ Xi and Xa Hi-C maps (clones merged). Scores represent B-B (Xi: S2-S2), B-A (Xi: S2-S1), and A-A (Xi: S1-S1) compartment interaction strength. DMSO and dTAG-13 treatment as in (B). (J) Cumulative distribution plots of allelic chromosome X and chromosome 13 expression by PRO-seq in RAD21-FKBP^degron^ lines. DMSO and dTAG-13 treatment as in (B).

The *Firre* locus also normally retains its cohesins on the Xi (**Fig.3E**, RAD21 track)(Bonora et al., 2018; Froberg et al., 2018; Giorgetti et al., 2016). It may thereby form a superloop with *Dxz4* through the retained cohesins, potentially through a cohesin-mediated loop extrusion mechanism across a 30Mb distance (Bansal et al., 2019; Bonora et al., 2018). Intriguingly, our allelic Hi-C map revealed a cohesin-dependent ‘stripe’ extending from *Dxz4* (**Fig.3D, inset**). This stripe (Vian et al., 2018) is indicative of a unidirectional loop extrusion whereby cohesin remains bound at the *Dxz4* anchor while extruding chromatin. Contrary to expectation, however, this stripe extended away from *Firre*, implying that *Dxz4* does indeed anchor loops, but loop extrusion may not be the mechanism by which it engages *Firre*. Indeed, the *Dxz4-Firre* superloop was not perturbed by cohesin depletion. It paradoxically became stronger with cohesin degradation (**Fig.3E, 4F**). Not only did the superloop strengthen on the Xi, it became visible on the Xa as well (**Fig.3E, 4F**). Thus, surprisingly, the *Dxz4*-*Firre* superloop is not cohesin-dependent and is in fact inhibited by cohesin.

We asked whether the superloop might depend instead on Polycomb, as the case for a subset of autosomal loops and the inter-*Hox* contacts (**Figs 1-2**). However, *Firre* and *Dxz4* were depleted for H3K27me3 relative to the rest of the Xi (**Fig.3E**), arguing against Polycomb-dependent looping for these loci. We noted that *Firre* and *Dxz4* belong to a group of “escapees”, X-linked genes that are not subject to XCI, and surmised that their escapee status might render them resistant to cohesin depletion. Intriguingly, allelic Hi-C analysis revealed a clustering of multiple escapee genes, including *Kdm5c*, *Ogt*, and *Bcor-Bcorl1* (**Fig.3F-G and 4F**). All such loops were reinforced by RAD21 degradation. While some escapees were previously observed to associate using 4C (Splinter et al., 2011), our high resolution allelic Hi-C maps enabled us to identify these superloops on a chromosome-wide basis and demonstrate that cohesin antagonizes their formation. No replicable changes to escapee expression was detected by PRO-seq at day 7 (8 hours degradation). A trend for decreased escapee expression could be detected by RNA-seq at day 10, but was only significant in one replicate **(Fig.4I).** We conclude that escapees form long-range loops through mechanisms other than cohesin-mediated loop extrusion and Polycomb-based co-clustering, and instead self-associate via active chromatin. We surmise that escapee clustering may aid in their persistent expression on the Xi, independently of cohesin and Polycomb.

### Transitory S1/S2 compartments on the Xi do not require cohesins

We next investigated effects on Xa and Xi compartments. Prior to XCI, the two X-chromosomes are organized into A/B compartments, much as any autosome. At the onset of XCI, Xist RNA — together with the PRC1 that it recruits — fuses the A/B compartments into larger intermediate structures called S1 and S2 compartments (**Fig.3A,** middle column) (Wang et al., 2019). Xist then recruits the non-canonical SMC protein, SMCHD1, to merge S1 and S2 compartments into a unique Xi-specific superstructure in which compartments are not visible (Wang et al., 2018). We asked how the transition from A/B to S1/S2 to the Xi superstructure is affected by cohesin perturbations. At days 7 and 10, S1/S2 compartments were visible on the control Xi as “checkerboard” patterns on a Pearson correlation heatmap, and also as alternating +/- segments in the 1st Principal Component of the Hi-C map (**Fig.3D,H,4D,G**), whereas classic A/B compartments were evident on the Xa (**Fig.3D,H,4E,G**). After 8 hours of cohesin degradation, these strength patterns were unchanged on both Xa and Xi (**Fig.3I, 4H**). Nor were there replicable, significant effects on Xi silencing on either day 7 or 10 (**Fig.3J, 4J**). While these findings are in line with the fact that up to 90% of cohesins are normally evicted from the Xi during XCI (Minajigi et al., 2015), the findings were somewhat unanticipated, given how much residual cohesin binding contributes to the Xi superstructure. Thus, like some Polycomb targets on autosomes, the Xi superstructure and transcriptomics appear to be resistant to cohesin loss. We surmise that its enrichment for Polycomb marks at least partially underlies the resistance.

### Forced cohesin retention disrupts the 3D structure of Xi and creates megadomains on Xa

We asked the opposite question: What would happen to the Xi if we forced cohesin retention? In our WAPL-FKBP^degron^ lines, 8 hours of WAPL depletion only succeeded in retaining cohesins at the 30Mb centromeric and distal telomeric ends of the Xi (**Fig.S4C**). We therefore focused analysis on 3 days of WAPL depletion (days 4-7), when cohesins became enriched 2-fold across the Xi (**Fig.5B**). A 2-fold enrichment also occurred on the Xa. Interestingly, although cohesins were 2-fold enriched on the Xi, its levels were not restored to Xa levels (**Fig.5B**). These findings suggest that recruitment of WAPL to the Xi may not be the sole means of cohesin eviction during XCI. Alternatively, it is possible that days 4-7 is too late a time point to fully restore cohesin levels. Nevertheless, the forced cohesin retention enabled us to examine consequences for Xi architecture and gene expression.

**Fig. 5.**
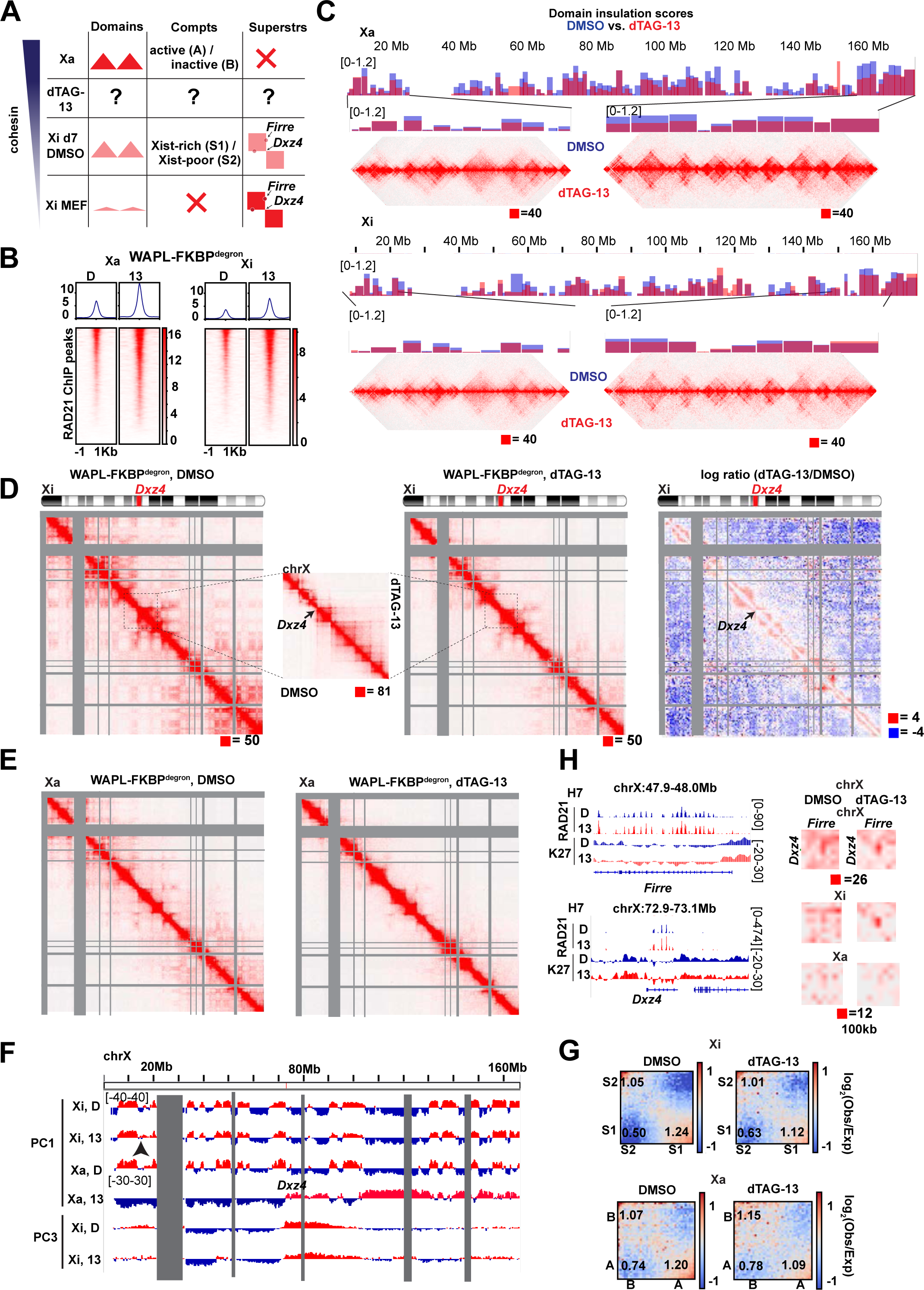
Cohesin retention results in increase of interactions spanning TAD borders and weakening of megadomains and compartments on the Xi. (A) Schematic of expected cohesin levels versus TAD, compartment, and superstructures detected on the X chromosome at different stages of XCI, based on prior studies. Levels corresponding to day 7 DMSO control and dTAG-13 treated cells indicated. (B) Metaplot and heatmaps of active (Xa) and inactive (Xi) X chromosome RAD21 binding by cChIP-seq in day 7 WAPL-FKBP^degron^ embryoid bodies treated for 3 days with 500nM dTAG-13 or DMSO. (C) TAD scores calculated from day 7 Xa and Xi WAPL-FKBP^degron^ allelic Hi-C interaction maps binned at 100kb. DMSO and dTAG-13 samples as in (B). Representative zoomed-in Hi-C maps shown for the indicated regions (50kb resolution, clones merged). (D) Chromosome-wide day 7 Xi WAPLFKBP^degron^ allelic Hi-C maps (250kb resolution, clones merged) and log2-ratio map (1Mb resolution). DMSO and dTAG-13 treatment as in (B). Inset (50kb resolution) showing ‘stripe’ at the megadomain border, *Dxz4*. (E) Chromosome-wide day 7 Xa WAPLFKBP^degron^ allelic Hi-C maps (250kb resolution, clones merged). DMSO and dTAG-13 treatment as in (B). (F) First and third Principal Component (PC1 and PC3) tracks of day 7 Xa and Xi WAPL-FKBP^degron^ Hi-C maps. DMSO and dTAG-13 treatment as in (B). (G) Saddle plot analysis of compartment strength in WAPL-FKBP^degron^ Xi and Xa Hi-C maps (clones merged). Scores represent B-B (Xi: S2-S2), B-A (Xi: S2-S1), and A-A (Xi: S1-S1) compartment interaction strength. DMSO and dTAG-13 treatment as in (B). (H) Zoomed-in day 7 composite X, Xi, and Xa Hi-C maps at *Dxz4-Firre* superloop (100kb resolution, clones merged). RAD21 and H3K27me3 cChIP-seq at superloop anchors shown for reference. DMSO and dTAG-13 treatment as in (B).

Intriguingly, forced cohesin retention did not substantially change strength of domains on the Xi (so-called “TADs”; **Fig.5C**) — contrasting with the marked attenuation seen when cohesins were depleted (**Fig.3C,4C**). Furthermore, the *Dxz4* megadomain border remained strong and the architectural stripe anchored at *Dxz4* appeared to lengthen (**Fig.5D**), consistent with cohesin retention and a unidirectional loop extrusion originating at *Dxz4*. The preservation of shorter-range looping interactions (**Fig.5C**) occurred at the expense of longer-range intra-megadomain interactions (**Fig.5D, right panel – subtractive map**). Longer-range interactions were also affected on the Xa (**Fig.5E**), in line with increased cohesin retention on the Xa as well. However, PC1 analysis showed that cohesin retention did not affect the appearance of transitory S1/S2 compartment structures except for disruptions at the distal centromeric end of the Xi (**Fig.5F, arrow**). Saddle plot analysis confirmed changes in compartments, with S1-S1 interactions weakening and S1-S2 interactions strengthening (**Fig.5G**), which could potentially explain the disruptions to S1/S2 compartments. Finally, no effects on the *Dxz4-Firre* superloop was observed (**Fig.5H**), further enforcing the idea that the superloop is cohesin-independent. Thus, forced cohesin retention resulted in failure to attenuate local looping interactions and disrupted the Xi superstructure without affecting superloops.

Findings for the Xa were equally intriguing. The Xa also enriched for cohesins by 2-fold after WAPL degradation (**Fig.5B**). While this retention did not affect topological domains (**Fig.5C**), A/B compartments could not be detected in PC1 of the Xa Hi-C map (**Fig.5F**). Instead, Xa adopted the megadomain-like pattern, which is typically only seen for the Xi (**Fig.5F**). The megadomain similarity between a cohesin-enriched Xa and the wildtype Xi could be seen in the 3^rd^ principle component of the Xi Hi-C map (PC3; **Fig.5F**). Saddle plot analysis showed weakened A-A interactions and strengthened A-B interactions (**Fig.5G**), which may partially account for the sudden change in the overall Xa structure. Thus, unexpectedly, megadomain organization can emerge independently of XCI.

### Failure of cohesin eviction compromises Xist spreading and Xi gene silencing

Previous work demonstrated that merging S1/S2 compartments is critical for Xist to spread and silence genes on the Xi (Wang et al., 2018). Given the changes in compartment structure (**Fig.5**), we asked if forced cohesin retention affected Xist spreading and Xi silencing. We performed CHART-seq to map genomic binding sites for Xist RNA in WAPL-depleted cells (dTAG-13 treated between days 4-7). Xist RNA still bound the Xi despite retained cohesins (**Fig.6A and S6A**). However, the localization appeared to be compromised, as there was a 2-fold drop in Xist coverage at the centromeric and telomeric ends (**Fig.6A,S6A,** Xist Δ track). Notably, these regions are furthest from the *Xist* locus. In the Hi-C contact map, these were also the regions that tend to lose contact with *Xist* (**Fig.6A,S6A,** top track). Statistical analysis confirmed Xist RNA depletion across the entire chromosome, with greatest effect size at the 30 Mb centromeric and telomeric ends (**Fig.6B, S6B**). Concomitantly, H3K27me3 levels decreased up to 4-fold at the Xi ends (**Fig.6A,S6A,** H3K27me3 tracks). Statistical analysis confirmed this decrease (**Fig.6C,S6C**). These effect sizes are Xi-specific in both biological replicates. While Xist and H3K27me3 coverages decreased significantly in both S1 and S2 compartments, the S1 compartment near the chromosomal ends were most affected (**Fig.6A, S6A**, yellow highlights) and the effect size was largest in S1 compartments overall (**Fig.S6F-G**). These data support the notion that weakened compartmentalization negatively impacts Xist and H3K27me3 spreading.

**Fig. 6.**
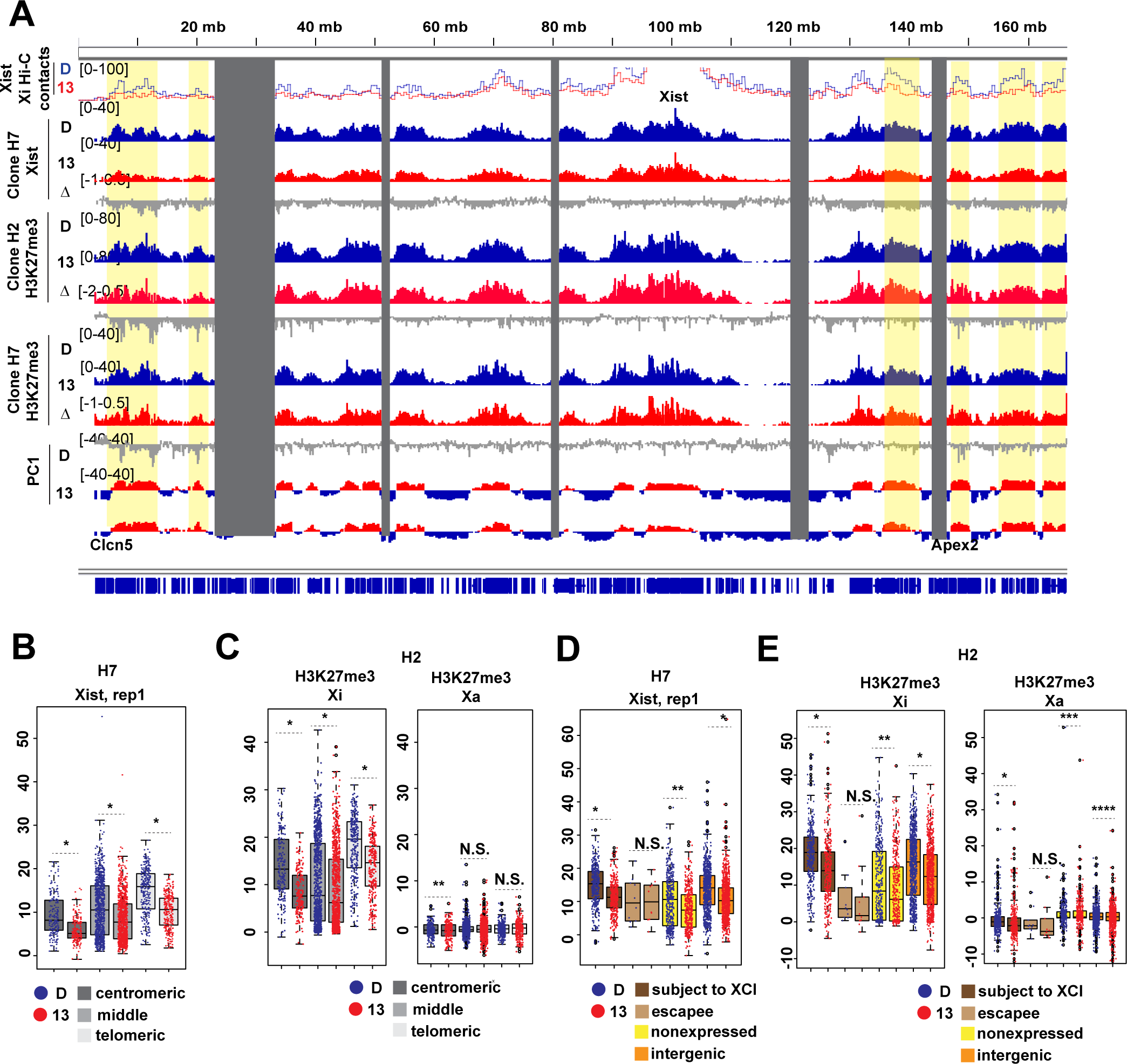
Retaining cohesin on the Xi early in differentiation leads to XCI defects, especially at the ends of the chromosome. (A) Tracks of Xi Hi-C contacts to the Xist transcription locus, Xist binding (CHART-seq) and H3K27me3 (cChIP-seq) in WAPL-FKBP^degron^ lines treated from days 4-7 of differentiation with DMSO (D) or 500nM dTAG-13 (13). Log fold ratio tracks (dTAG-13/DMSO) are also shown (Δ). Principal component corresponding to compartments on the Xi (PC1, positive S1, negative S2) shown for reference. (B) Boxplot showing Xist coverage over 100kb bins in the centromeric most 30Mb, middle 100Mb, and telomeric most 30Mb of the X chromosome in WAPL-FKBP^degron^, clone H7 treated as in (A). Significant p-values by Wilcoxon signed rank test indicated by * (p<2.2e-16). (C) Boxplots showing allelic H3K27me3 coverage over 100kb bins in the centromeric most 30Mb, middle 100Mb, and telomeric most 30Mb of the X chromosome in WAPL-FKBP^degron^, clone H2 treated as in (A). Significant p-values by Wilcoxon signed rank test indicated by * (p<2.2e-16) and ** (p<0.001187). N.S., not significant. (D) Boxplot showing average Xist density over different classes of genes on the X chromosome in WAPL-FKBP^degron^, clone H7 treated as in (A). Significant p-values by Wilcoxon signed rank test indicated by * (p<2.2e-16). N.S., not significant. (E) Boxplot showing average allelic H3K27me3 density over different classes of genes on the X chromosome in WAPL-FKBP^degron^, clone H2 treated as in (A). Significant p-values by Wilcoxon signed rank test indicated by * (p<2.2e-16), ** (p<8.094e-16), *** (p<0.00206), **** (p< 0.0001193). N.S., not significant.

It is possible that the increased number of longer chromatin loops (**Fig.1K,M**) also plays a role in weakened Xist and PRC2 spreading. As looping contacts mainly increased in the 1-10Mb range (**Fig.1K**), we examined how spreading was affected within loops of <10Mb. While Xist coverage decreased within 10Mb, H3K27me3 was not replicably changed (**Fig.S6H**). Therefore, while additional chromatin loops may inhibit Xist spreading, this does not seem to be sufficient to impede H3K27me3 deposition at short ranges.

Finally, we investigated effects on Xi gene silencing. Xist and H3K27me3 coverages were significantly decreased over genes subject to XCI (**Fig.6D-E,S6D-E**). They were also reduced over intergenic regions, but not over escapees which normally lacked Xist and H3K27me3. We performed allele-specific RNA-seq analysis in WAPL-depleted cells (dTAG-13 treated days 4-7) and observed a significant cumulative shift in expression from the Xi (**Fig.7A**) — indicating either a failure of inactivation during the establishment phase, or an initial inactivation and then a partial reactivation in later stages. Consistent with compromised spreading of the Xist-PRC2 complex, the failure of silencing was greater at the centromeric and telomeric 30Mb ends (**Fig.7B**). Examination of specific loci revealed strong (>1.7-fold) upregulation from various genes, including *Clcn5*, *Apex2*, *Tspan7*, *Spin2c*, and *Ap1s2* (**Fig.7C**) — all of where were located at the chromosome ends. We conclude that aberrant cohesin retention disrupts the ability of the Xist-PRC2 complex to spread at long range to the ends of the Xi, with consequences for gene silencing.

**Fig.7.**
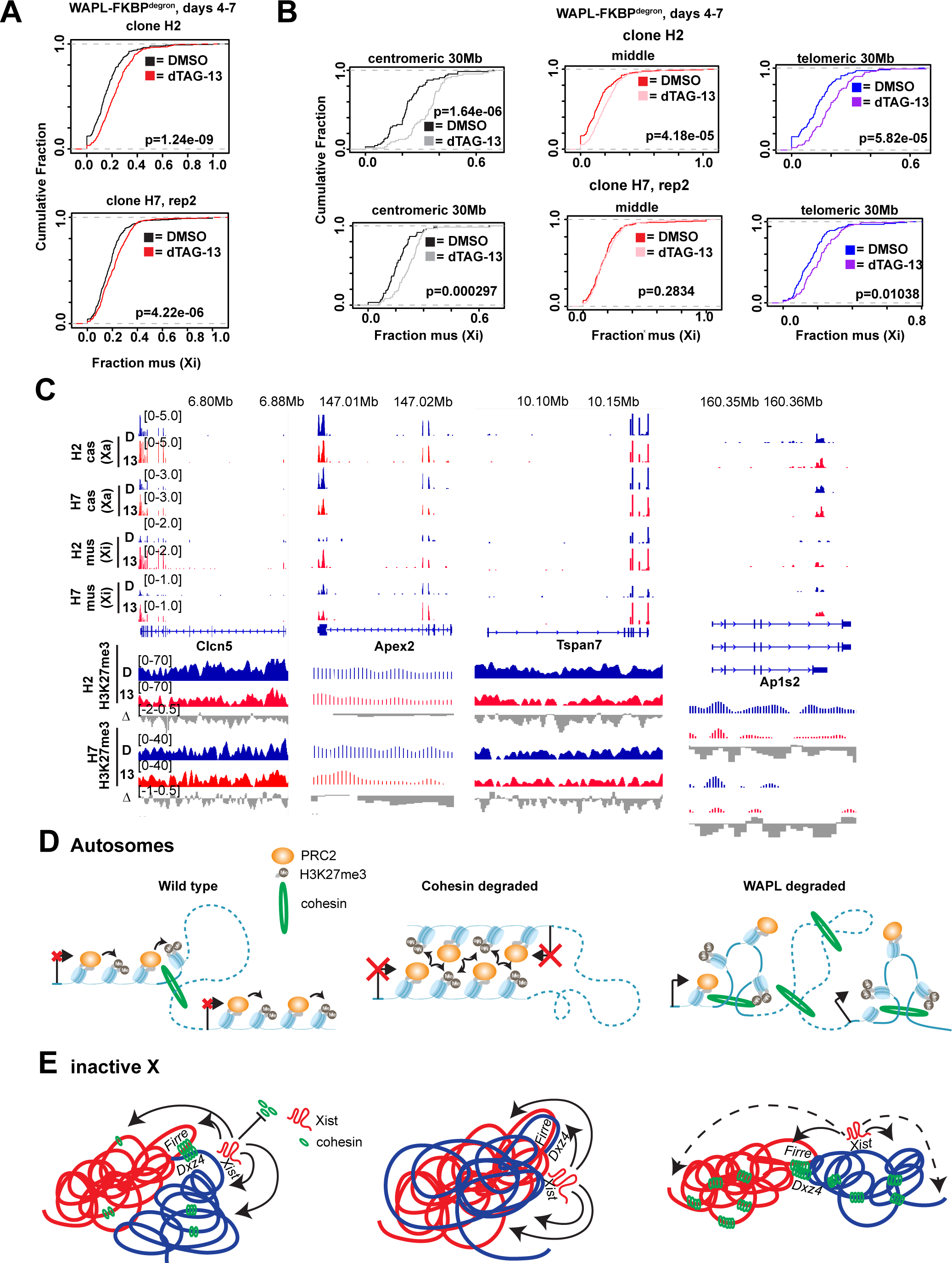
Selective cohesin eviction and retention are required to maintain Polycomb targets, fold the Xi, and properly silence Xi genes. (A) Cumulative distribution plots of chromosome-wide Xi expression in 2 WAPL-FKBP^degron^ clones treated from days 4-7 of differentiation with DMSO or 500nM dTAG-13. P-value calculated by the KS test. (B) Cumulative distribution plots of Xi expression from the centromeric most 30Mb, middle 100Mb, and telomeric most 30Mb of the chromosome in 2 in 2 WAPL-FKBP^degron^ clones treated as in (A). P-value calculated by the KS test. (C) Examples of genes at the far centromeric (Clcn5,Tspan7) and telomeric (Apex2,Ap1s2) ends of the X chromosome. Allelic RNA-seq and H3K27me3 cChIP-seq shown for 2 WAPL-FKBP^degron^ clones treated as in (A). (D) Model: cohesin tempering of Polycomb targets on autosomes. (E) Model: cohesin tempering of Xist and Polycomb spreading on the Xi.

## DISCUSSION

Large and finer scale 3D genome structures have long been hypothesized to play a pivotal role in gene regulation (de Wit et al., 2015; Guo et al., 2015; Merkenschlager and Nora, 2016). However, recent degron studies targeting architectural factors and their genomic binding sites have challenged this model in that very few genes are dysregulated as a consequence. Here, our study revisited the genomic structure-function relationship through the lens of XCI and demonstrate that a fine balance between cohesin eviction and retention is necessary to shape the Xi. Additionally, this balance is required to protect against inappropriate inter-chromosomal contacts and to temper Polycomb-mediated gene repression on autosomes.

While it is now established that the Xist RNA evicts cohesins and the Xi is generally depleted of these architectural factors, it was previously not known whether some cohesins must be retained in order for proper architectural transformation into a silent chromosome. By toggling cohesin levels up or down at will, our present work demonstrates that XCI requires that cohesins be evicted from some regions and retained on others. For instance, the sharp megadomain border and intra-megadomain interactions require cohesin. Depleting cohesin ablates the megadomain border, whereas forcing cohesin retention does not compromise the sharp border but reduces intra-megadomain interactions. Likewise, the finer-scale loop domains or “TADs” on the Xi require cohesin. When cohesin is depleted, these structures disappear almost entirely.

Our study also demonstrates that Xi compartments — specifically the S1/S2 compartments — are weakened by stabilization of cohesin. Superloops, such as the those between *Dxz4-Firre* and between escapees *Kdm5c* and *Ogt*, are also antagonized by cohesins and strengthened when cohesins are degraded. Thus, the Xi indeed does not become stripped of all cohesin and selective regions must retain cohesin binding in order to construct the Xi superstructure. An interesting question for future consideration is how Xist evicts one set of cohesins while retaining others on the Xi.

The present work also reveals that Xist and Polycomb spreading and gene silencing all depend on cohesin eviction across the Xi. Forced cohesin retention results in reduced spreading of Xist RNA and H3K27me3 enrichment across the whole Xi, but especially at the distal ends. Gene silencing was also blunted to varying degrees along the whole chromosome, with the strongest effects again at the Xist-distal ends. Expression of escapee genes, by contrast, is not affected by cohesin retention. However, cohesin depletion led to a surprising reinforcement of clustering of various escapees. The *Dxz4-Firre* superloop also strengthened upon cohesin depletion. It is possible that the depletion of cohesin that normally occurs on the Xi reinforces clustering of escapees to ensure their continued expression. Intriguingly, a recent study implicated *Firre* RNA as a regulator of the Xi and autosomes *in trans* (Fang et al., 2020). Future work will be needed to determine if interactions of the *Firre* genomic locus could be involved as well.

Finally, our data show that the effects of cohesin imbalances are not confined to the Xi. Indeed, megadomains appear mysteriously on the Xa when cohesins are aberrantly enriched, suggesting that enhanced local contacts may spontaneously drive partitioning of an X-chromosome into two large domains. Importantly, this means that megadomain organization can emerge independently of Xist and XCI. A *Dxz4-Firre* interaction also appeared on the Xa after cohesin degradation, also suggesting that the superloop could be a consequence of cohesin depletion rather than XCI. On autosomes, cohesins antagonize interactions between Polycomb targets and temper Polycomb-mediated repression (Rhodes et al., 2020). Unexpectedly, the effects on Polycomb domains are not limited in cis, as loss of cohesin strengthens clustering of Polycomb targets *in trans,* as exemplified by trans-interactions among the four *Hox* gene clusters. Inter-chromosomal bridges between the *Hox* loci are especially prominent during differentiation, in contrast to mESCs, where such interactions are not detected in control conditions.

Our data also indicate that cohesins play an important role in Polycomb-mediated gene repression. When cohesin is ablated, trans-interactions strengthen between chromosomes. When cohesins are aberrantly retained, such interactions are disrupted and expression of the Polycomb targets are dramatically upregulated. Notably, while autosomal gene expression in differentiating ES cells is not hugely affected by cohesin imbalance in the short term, Polycomb targets are particularly dysregulated by either removing or stabilizing cohesin. These findings may cast light on why this and previous RAD21 degron studies detected relatively little gene dysregulation— as cohesin-independent loops may contribute to maintaining proper gene expression in these cases.

Altogether, the relationship between cohesin and gene expression on autosomes is analogous to that observed on the Xi (**Fig.7D-E**). On autosomes under wild type conditions, current models postulate that cohesin translocates along chromatin during the process of loop extrusion, passing through Polycomb domains (**Fig.7D**, left). Polycomb targets participating in interactions with each other tend to be highly repressed and are therefore indifferent to the presence of cohesin. However, certain Polycomb targets which are not engaged in interactions can become further repressed upon loss of cohesin. As Polycomb-anchored loops appear to be reinforced through cohesin loss, it is possible that this occurs due to increased interactions between previously non-interacting regions (**Fig.7D**, middle). As the Xi as a whole is highly repressed and covered with Polycomb that forms self-associating compartments, it is logical that Xi expression would also be indifferent to the loss of cohesin, similar to interacting Polycomb targets on autosomes (**Fig.7E**, middle). When WAPL is degraded, cohesin levels increase genome-wide, weakening interactions between Polycomb domains. Such disruptions lead to upregulation of Polycomb targets previously engaged in such interactions (**Fig.7D**, right). We also observe upregulation across the Xi. In contrast to autosomes, repression on the Xi is mediated by Xist (**Fig.7E**, left). Therefore, we observe the largest impact on gene silencing at the distal ends of the chromosome furthest from Xist, where the most Hi-C contacts are lost as well (**Fig.7E**, right). In conclusion, we propose that a key role of cohesins is to organize local chromatin and prevent ectopic long-range cis- and trans-interactions. In doing so, cohesins tempers repression of Polycomb targets as well as X chromosome inactivation.

## AUTHOR CONTRIBUTIONS

A.J.K. and J.T.L. conceived the project, analyzed data, and wrote the paper. A.J.K. performed all bioinformatics analyses. D.C. and H.S. performed ChIP-seq in parental TST mESCs. A.J.K. performed all other experiments. B.N. generated dTAG-13 and provided advice in generation of FKBP^degron^ lines.

## ACKNOWLEDGEMENTS

We thank J. Froberg and C.-Y. Wang for critical reading of the manuscript, and we thank all lab members for support. We thank N. Gray for support and the generous gift of dTAG-13. The authors would like to thank the Nascent Transcriptomics Core at Harvard Medical School, Boston, MA for performing PRO-seq library construction. Nucleosomes in Fig.7 were created with BioRender.com. A.J.K. is supported by 1F31HD100109-01 and J.T.L. by grants from the National Institutes of Health (9R01HD097665-09) and the Howard Hughes Medical Institute. B.N. is supported by an American Cancer Society Postdoctoral Fellowship (PF-17-010-01-CDD) and the Katherine L. and Steven C. Pinard Research Fund.

## SUPPLEMENTAL FIG.LEGENDS

**Fig. S1.**
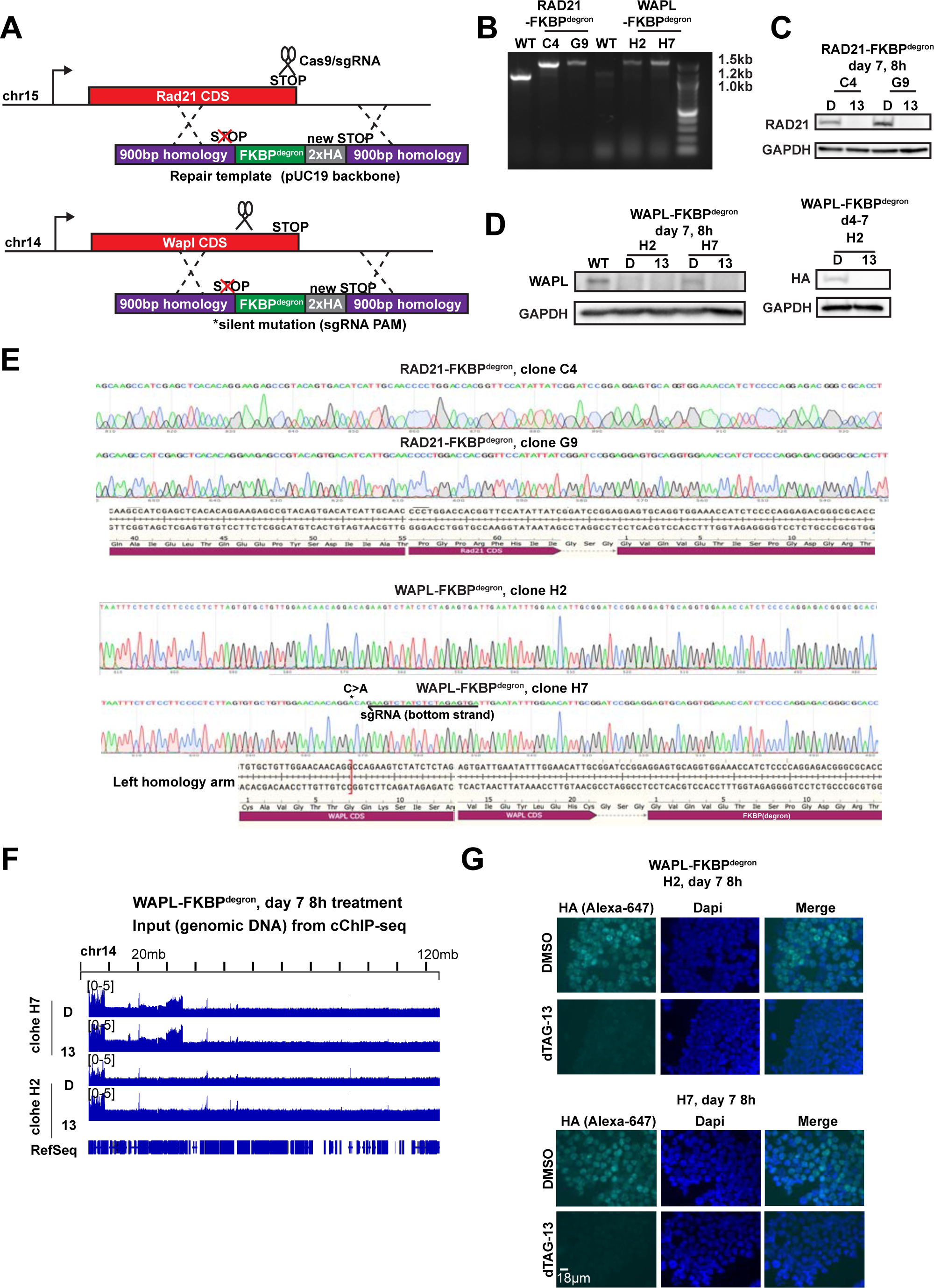
Generation of FKBP^degron^ lines. (Related to Fig.1) (A) Outline of protocol used to generate RAD21 and WAPL-FKBP^degron^ mESCs (B) PCR amplifying knock-in site of parental TST mESC lines (WT) and RAD21 and WAPL-FKBP^degron^ clones. One PCR primer was located outside of the homology arms to ensure that only the endogenous locus was amplified. FKBP^degron^ tag is ∼400bp. (C) Western blots for RAD21 and GADPH loading control in RAD21-FKBP^degron^ lines at day 7 of differentiation. Samples treated with DMSO vehicle control (D) or 500nM dTAG-13 (13). (D) Western blots for WAPL and GADPH loading control in WAPL-FKBP^degron^ lines at day 7 of differentiation. Samples treated with DMSO vehicle control (D) or 500nM dTAG-13 (13). (E) Sanger sequencing of RAD21-FKBP^degron^ and WAPL-FKBP^degron^ lines. (F) Comparison of genomic DNA (input DNA) in WAPL-FKBP^degron^ lines at chromosome 14. Sharp dip in WAPL-FKBP^degron^ , clone H7 input corresponds to the end of the right 900bp homology arm used for knock-in of the tag. Reads mapping to the plasmid backbone used for the homology repair template were also found here, although they did not appear to be incorporated into the main Wapl transcript. (G) Immunofluorescence of WAPL-HA in WAPL-FKBP^degron^ lines at day 7 of differentiation. Samples treated with DMSO vehicle control (D) or 500nM dTAG-13 (13).

**Fig. S2.**
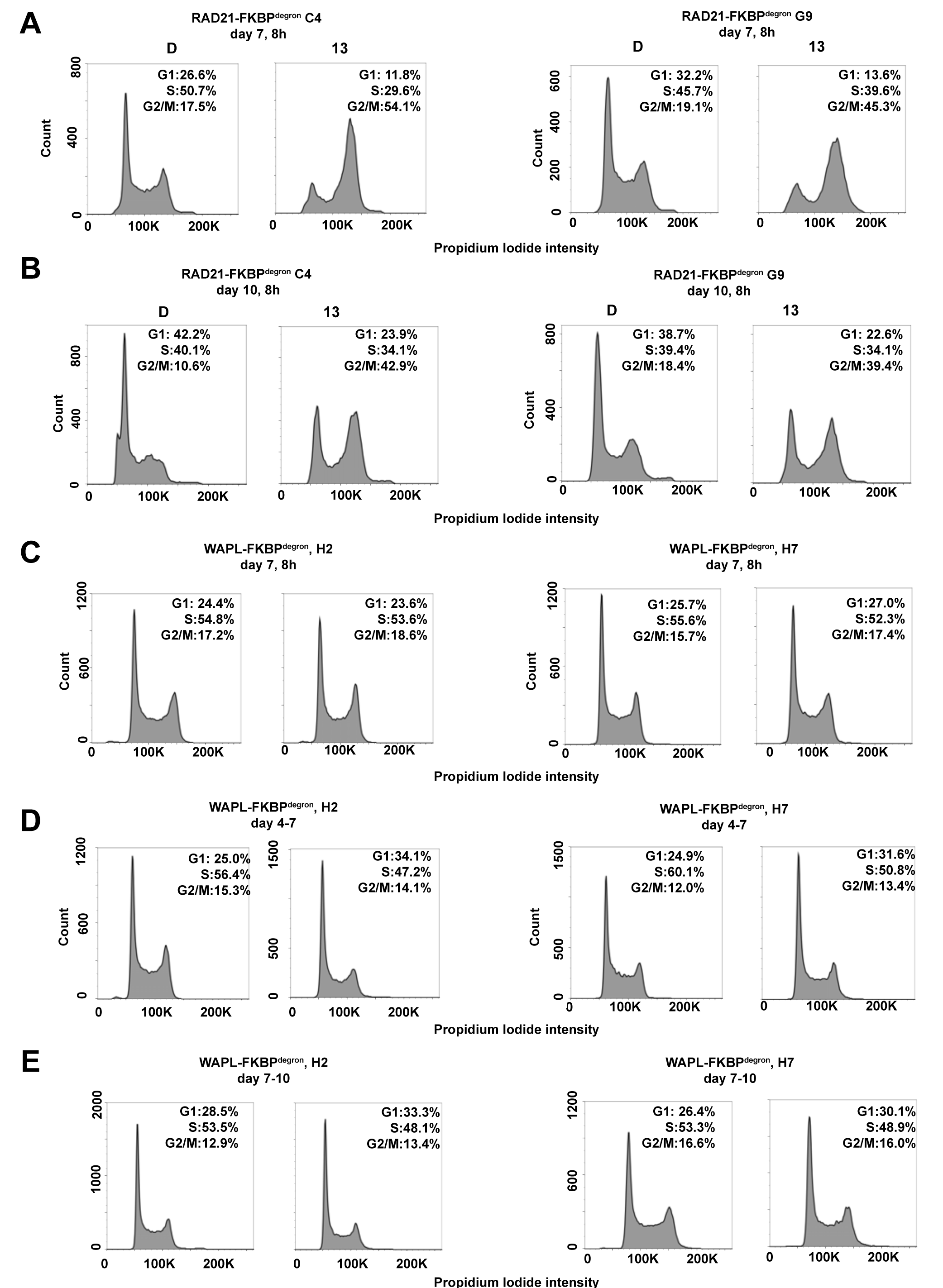
Cell cycle profiling of FKBP^degron^ lines. (Related to Fig.1) (A) DNA content profiles via propidium iodide staining of RAD21-FKBP^degron^ lines at day 7 of differentiation. Samples treated with DMSO vehicle control (D) or 500nM dTAG-13 (13) as indicated. (B) DNA content profiles via propidium iodide staining of RAD21-FKBP^degron^ lines at day 10 of differentiation. (C) DNA content profiles via propidium iodide staining of WAPL-FKBP^degron^ lines at day 7 of differentiation. (D) DNA content profiles via propidium iodide staining of WAPL-FKBP^degron^ lines at day 7 of differentiation. (E) DNA content profiles via propidium iodide staining of WAPL-FKBP^degron^ lines at day 10 of differentiation.

**Fig. S3.**
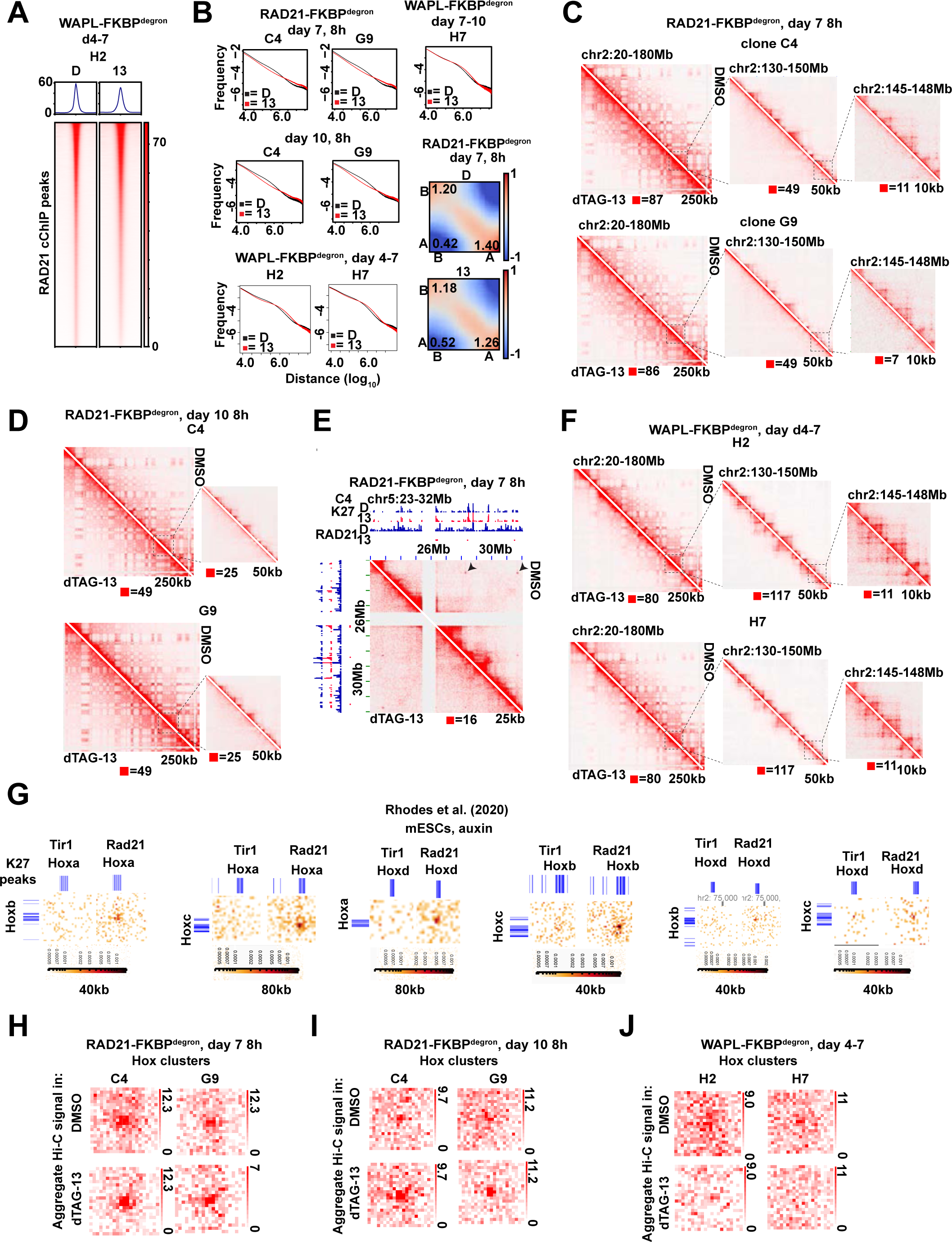
Replication of architectural changes across FKBP^degron^ clones. (Related to Fig. s 1 and 2) (A) Metaplot and heatmap of RAD21 binding by cChIP-seq in WAPL-FKBP^degron^ clone H2. (B) Frequency versus distance plots for Hi-C interaction maps across individual FKBP^degron^ clones at all time points used in this study. Samples treated with DMSO vehicle control 500nM dTAG-13 as indicated. Saddle plot analysis of compartment strength in RAD21-FKBP^degron^ Hi-C maps at day 7 (clones merged) also shown. Scores represent B-B, B-A, and A-A compartment interaction strength.Representative sections of day 7 Hi-C maps at different resolutions for RAD21-FKBP^degron^ clones. (C) Representative sections of day 10 Hi-C maps at different resolutions for RAD21-FKBP^degron^ clones. (D) Section of day 7 RAD21-FKBP^degron^ Hi-C maps (clones merged) displaying example of cohesin-independent loops not anchored by H3K27me3 (arrows). (E) Representative sections of day 7 Hi-C maps at different resolutions for WAPL-FKBP^degron^ clones. (F) Aggregate Peak Analysis plots of aggregate RAD21-FKBP^degron^ Hi-C signal at day 7 (clones merged) over regions anchored by Hox clusters. (G) Aggregate Peak Analysis plots of aggregate RAD21-FKBP^degron^ Hi-C signal at day 10 (clones merged) over regions anchored by Hox clusters. (H) Aggregate Peak Analysis plots of aggregate WAPL-FKBP^degron^ Hi-C signal at day 7 (clones merged) over regions anchored by Hox clusters.

**Fig. S4.**
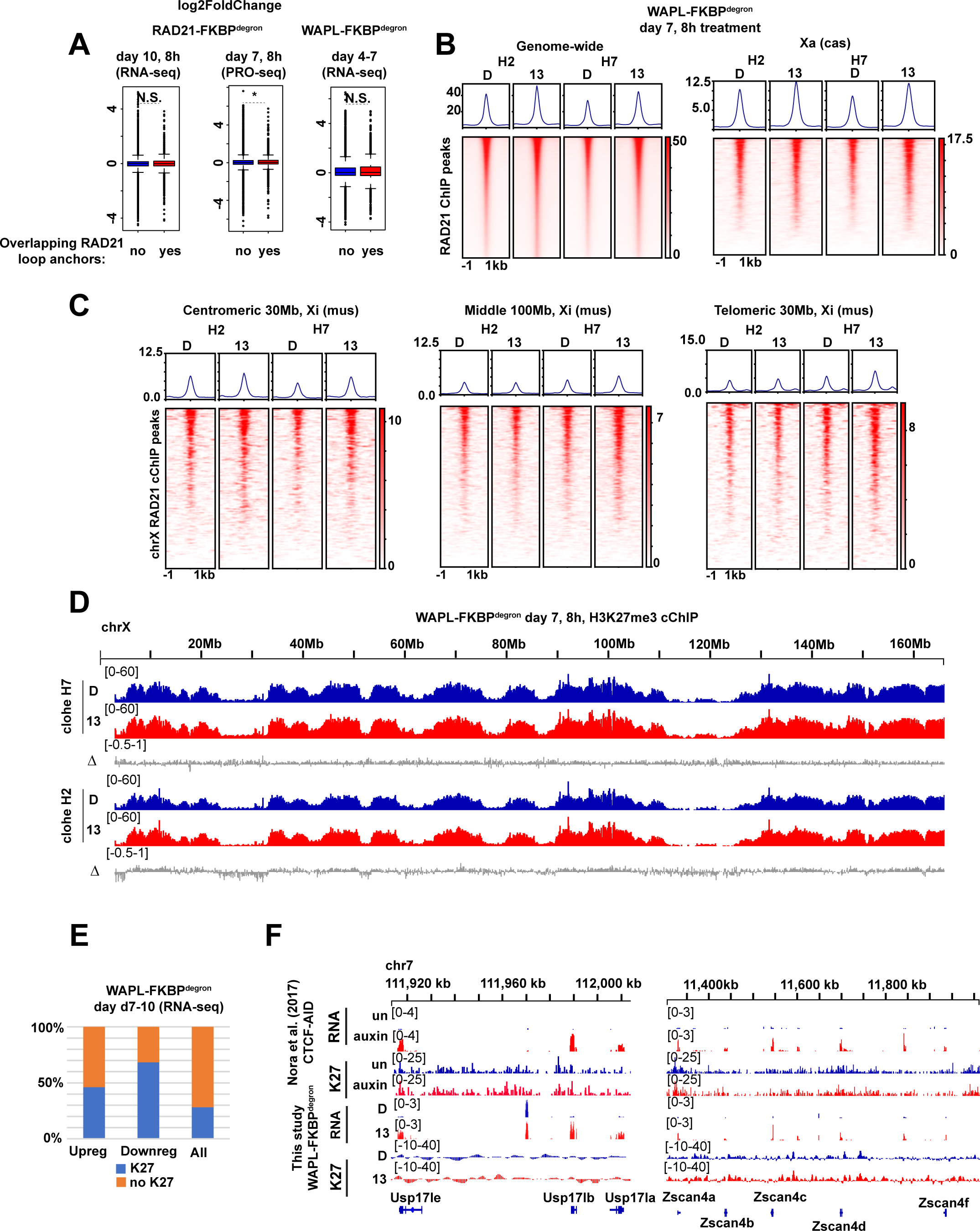
Additional analysis of cChIP after short term WAPL degradation and gene expression changes after 3 day degradation. (Related to Fig.3) (A) Log2-fold (dTAG-13/DMSO) change in FKBP^degron^ lines of genes overlapping RAD21-bound anchors of loops. Loops called in respective DMSO control samples. Significant p-values by Wilcoxon rank test indicated by * (p<2.772e-06). N.S., not significant. (B) Metaplot and heatmaps of RAD21 binding by H3K27me3 cChIP-seq after 8 hours of WAPL degradation genome-wide (left) and on the active X chromosome (right). (C) Metaplot and heatmaps of RAD21 binding by H3K27me3 cChIP after 8 hours of WAPL on the centromeric 30Mb, middle 100Mb, and telomeric 30Mb of the inactive X chromosome. (D) Tracks of H3K27me3 by cChIP-seq on the X chromosome after 8 hours of WAPL degradation or DMSO control treatment. Log fold ratio tracks (dTAG-13/DMSO) are also shown (Δ). (E) Genes upregulated or downregulated upon WAPL degradation from days 7-10 (relative to DMSO control) overlapping with H3K27me3 cChIP-seq peaks. All genes shown for reference. (F) RNA-seq and H3K27me3 ChIP-seq over the *Usp17* and *Zscan4* gene clusters in CTCF-AID degron lines after 4 days of auxin treatment or untreated cells (Nora et al., 2017) or after 3 days of DMSO or dTAG-13 treatment (500nM) in WAPL-FKBP^degron^ cells early in differentiation (days 4-7).

**Fig. S5.**
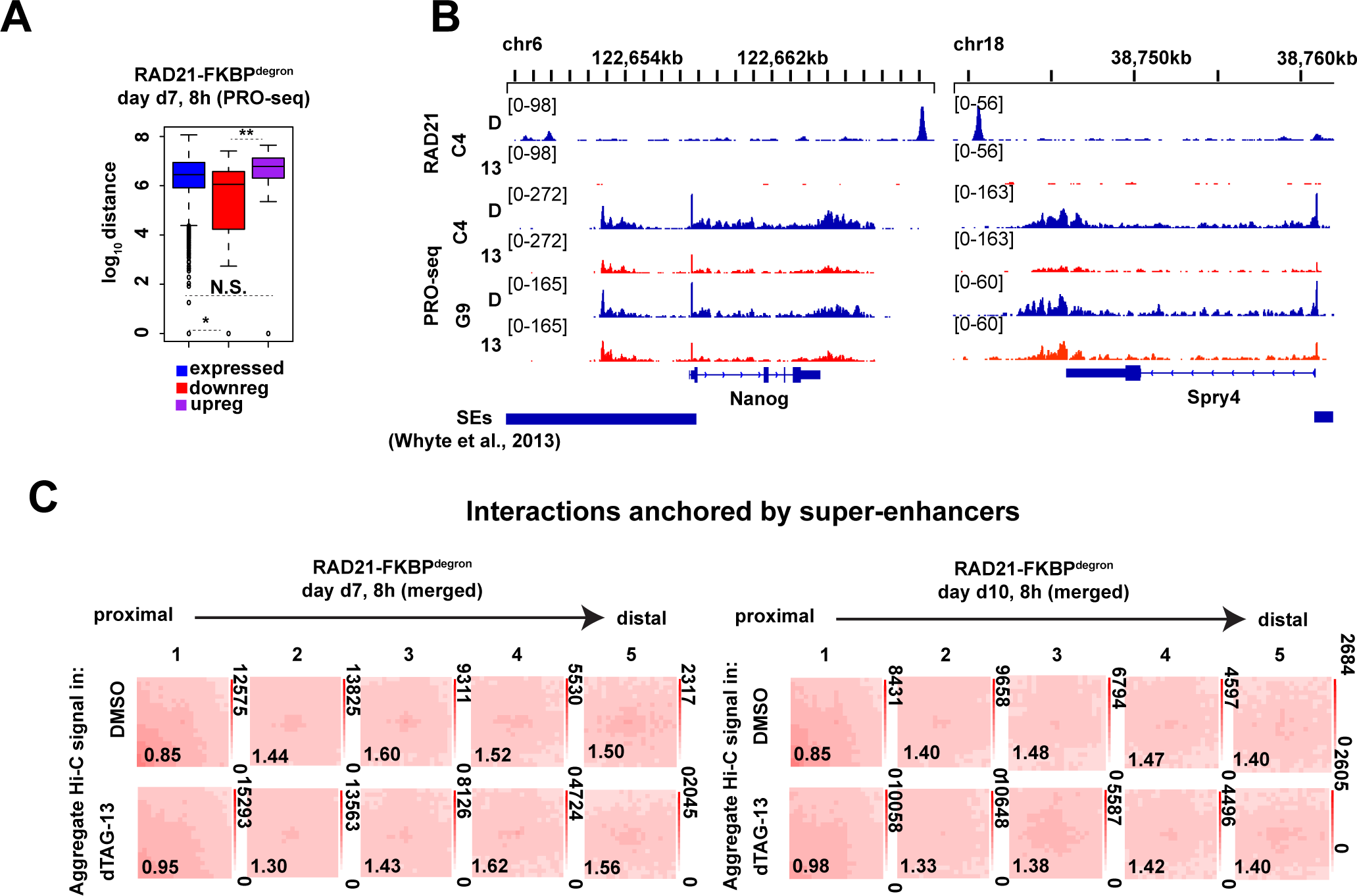
Cohesin is required for proper folding of the inactive X chromosome at day 10 of differentiation (Related to Fig.4) (A) Schematic of expected cohesin levels versus TAD, compartment, and superstructures detected on the X chromosome at different stages of XCI, based on prior studies. Levels corresponding to DMSO control and dTAG-13 treated cells indicated. (B) Metaplot and heatmaps of active (Xa) and inactive (Xi) X chromosome RAD21 binding by uncalibrated ChIP-seq in day 10 RAD21-FKBP^degron^ embryoid bodies treated for 8 hours with 500nM dTAG-13 or DMSO. (C) TAD scores calculated from day 10 Xa and Xi RAD21-FKBP^degron^ allelic Hi-C interaction maps binned at 100kb. DMSO and dTAG-13 samples as in (B). Representative zoomed-in Hi-C maps shown for the indicated regions (50kb resolution, clones merged). (D) Chromosome-wide day 10 Xi RAD21-FKBP^degron^ Hi-C maps (250kb resolution, clones merged) and log2-ratio map (1Mb resolution). DMSO and dTAG-13 treatment as in (B). (E) Chromosome-wide day 10 Xa RAD21-FKBP^degron^ Hi-C maps (250kb resolution, clones merged) and log2-ratio map (1Mb resolution). DMSO and dTAG-13 treatment as in (B). (F) Zoomed-in day 7 composite X, Xi, and Xa Hi-C maps at *Dxz4-Firre* superloop (100kb resolution, clones merged), *Kdm5c-Ogt* loop (250kb resolution, clones merged) and *Bcor-Bcorl1* loop (100kb resolution, clones merged). RAD21 and H3K27me3 cChIP-seq at superloop anchors shown for reference. DMSO and dTAG-13 treatment as in (B). (G) First Principal Component (PC1) tracks of day 7 Xa and Xi RAD21-FKBP^degron^ Hi-C maps. DMSO and dTAG-13 treatment as in (B). (H) Saddle plot analysis of compartment strength in RAD21-FKBP^degron^ Xi and Xa Hi-C maps (clones merged). Scores represent B-B (Xi: S2-S2), B-A (Xi: S2-S1), and A-A (Xi: S1-S1) compartment interaction strength. DMSO and dTAG-13 treatment as in (B). (I) Xi escapee (including *Ogt, Bcor1, Bcorl1*) gene expression in RAD21-FKBP^degron^ lines. Significant p-values by Wilcoxon signed rank test indicated by * (p<0.001663), ** (p<0.00232), *** (p<0.0002441). N.S., not significant. (J) Cumulative distribution plots of allelic chromosome X and chromosome 13 expression by RNA-seq in RAD21-FKBP^degron^ lines. DMSO and dTAG-13 treatment as in (B).

**Fig. S6.**
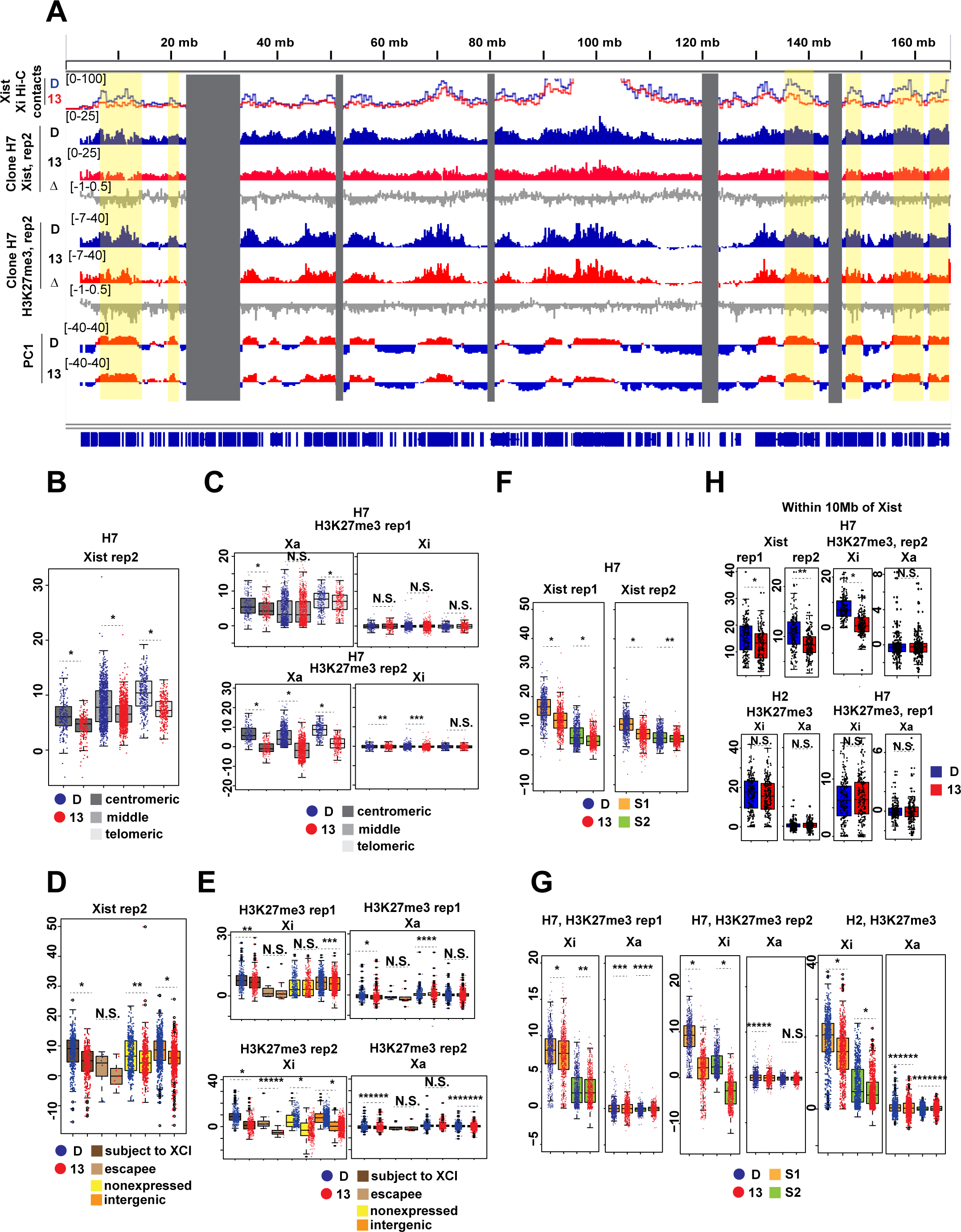
Additional biological replicate for Xist CHART-seq and H3K27me3 cChIP-seq to examine spreading defects. (Related to Fig.6). (A) Tracks of CHART-seq (Xist) and H3K27me3 cChIP-seq over the X chromosome and allelic RAD21 cChIP over the Xa and Xi in WAPL-FKBP^degron^ lines treated from days 4-7 of differentiation with DMSO (D) or 500nM dTAG-13 (13). Log fold ratio tracks (dTAG-13/DMSO) are also shown (Δ). Principal component corresponding to compartments on the Xi in same samples shown for reference (PC1). Hi-C contacts to the 1Mb bin containing the Xist transcription locus in same conditions also shown. (B) Boxplot showing Xist coverage over 100kb bins in the centromeric most 30Mb, middle 100Mb, and telomeric most 30Mb of the X chromosome in WAPL-FKBP^degron^ lines treated as in (A). Significant p-values by Wilcoxon signed rank test indicated by * (p<2.2e-16). (C) Boxplots showing allelic H3K27me3 coverage over 100kb bins in the centromeric most 30Mb, middle 100Mb, and telomeric most 30Mb of the X chromosome in WAPL-FKBP^degron^ lines treated as in (A). Significant p-values by Wilcoxon signed rank test indicated by * (p<2.2e-16), ** (p<0.03), and *** (p<2.2e-05). N.S., not significant. (D) Boxplot showing average Xist density over different classes of genes on the X chromosome in WAPL-FKBP^degron^ lines treated as in (A). Significant p-values by Wilcoxon signed rank test indicated by * (p<2.2e-16) and ** (p<1.049e-14). N.S., not significant. (E) Boxplot showing average allelic H3K27me3 density over different classes of genes on the X chromosome in WAPL-FKBP^degron^ lines treated as in (A). Significant p-values by Wilcoxon signed rank test indicated by * (p< 2.2e-16), ** (p< 2.79e-15), *** (p< 9.04e-13), **** (p< 0.001573), ***** (p< 0.001954), ****** (p< 2.239e-05), ******* (p< 1.308e-05). N.S., not significant. (F) Boxplot showing Xist average density over 100kb bins in S1 and S2 compartments. Significant p-values by Wilcoxon signed rank test indicated by * (p<2.2e-16) and ** (p<4.13e-09). (G) Boxplot showing average allelic H3K27me3 density over 100kb bins in S1 and S2 compartments. Significant p-values by Wilcoxon signed rank test indicated by * (p<2.2e-16), **(p<0.0082), *** (p< 0.005), ****(p<1.86e-10), ***** (p<1.89e-08), ****** (p<2.127e-05), ******* (p<0.0008859). (H) Boxplots showing average Xist and allelic H3K27me3 density over 100kb within 10Mb of the Xist transcription locus. Significant p-values by Wilcoxon signed rank test indicated by * (p<1.543e-05), **(p<4.026e-13).

## MATERIALS AND METHODS

### Cell Lines

RAD21-FKBP^degron^ and WAPL-FKBP^degron^ mESCs were generated by transfecting *Tsix^TST^*^/+^ mESCs with (1) a plasmid (pSpCas9-(BB) 2A Puro) expressing Cas9 with an sgRNA targeted to the STOP codon (in the case of Rad21) or to a region in the last codon near the STOP codon (in the case of Wapl) (2) a homology repair template plasmid including 900bp homology arms around the STOP codon, the FKBP^degron^ tag in a pUC19 backbone and (3) a homology-directed repair plasmid (pMB 1609-pRR-EGFP) with the sgRNA target site cloned in. Specifically, 500,000 TST mESCs were transfected using Lipofectamine 3000 1.5ug of pSpCas9-(BB) 2A Puro, 1.5ug of the non-linearized repair plasmid, and 400ng of pMB 1609-pRR-EGFP. Transfected cells were GFP sorted after 48 hours and the entire GFP positive pool plated. After 5 days, the pool was split and plated at low density, and after 5 more days, clones were picked from the low-density plate in a 96-well plate. Clones were grown to confluency, then triplicate plated in 96-well plates. One 96-well plate was used to PCR screen clones. Clones were then thawed from one of the triplicate plates, expanded, and re-screened by PCR and Sanger sequenced to confirm correct knock-in of the FKBP^degron^ tag.

### ES cell differentiation and induced degradation

Undifferentiated mESCs were grown on γ-irradiated MEF feeders in medium containing DMEM, high glucose, GlutaMAX supplement, pyruvate, 15% Hyclone FBS (Sigma), 25 mM HEPES pH 7.2-7.5, 1x MEM non-essential amino acids, 1x Pen/Strep, 0.1 mM βME, and 500 U/mL ESGRO recombinant mouse Leukemia Inhibitory Factor (LIF) protein (Sigma, ESG1107) at 37°C with 5% CO2. To induce differentiation in EBs, mESCs were trypsinized and resuspended in the same media without LIF. For the first 3 days of differentiation, EBs were grown in suspension. On day 4 and subsequent days, EBs were grown on gelatinized plates until harvested at day 7 or day 10.

Degradation of RAD21 or WAPL in FKBP^degron^ lines during differentiation was induced by adding a 1:20,000 dilution of 10mM dTAG-13 in DMSO to cell culture media (500nM final concentration) for the indicated time frames. The small molecule dTAG-13 was synthesized as previously described (Erb et al., 2017; Nabet et al., 2018). For DMSO vehicle controls, the same volume of DMSO alone was added to cell culture media.

### Western blot

Cells were counted and ∼100,000 cells per sample lysed in 100ul 1X Laemmli buffer. Protein extracts were boiled for 5 min at 95C. Equal cell counts (5-15ul) were loaded and resolved by SDS page. Protein was transferred to an Immobilon-P PVDF membrane (EMD Millipore). After transfer, membrane was blocked with 5% milk in PBST (PBS/0.05% Tween-20) for 30 min at room temperature. Membrane was incubated with primary antibody (1:1000 for RAD21 and WAPL, 1:20,000 for GAPDH) in 5% milk in PBST at 4C overnight, then washed three times with PBST. Membrane was then blocked with secondary antibody-HRP conjugate (1:5000) in 5% milk in PBST for 1 hour at room temperature. Membrane was incubated with Western Lightning Plus-ECL (PerkinElmer) and exposed using the Bio-Rad ChemiDoc MP Imaging System.

### Immunofluorescence

Immunofluorescence was performed as described previously with minor modifications (Del Rosario et al., 2019). Briefly, cells on glass slides were first permeabilized 2 min on ice with CSKT (100 mM NaCl, 300 mM Sucrose, 10 mM PIPES pH 6.8, 3 mM MgCl2, 0.5% Triton X-100) and then fixed in PBS 4% paraformaldehyde for 10 min at room temperature followed by a PBS wash step. Fixed cells were blocked (PBS, 0.2% Tween-20, 4% bovine serum albumin (BSA), 2% normal goat serum (NGS) (Sigma)) for 30 min at room temperature and incubated with primary antibody in a humid chamber at 4C overnight. Antibodies were diluted in blocking buffer. Samples were washed with PBS300.2T (PBS supplemented to 300 mM NaCl, 0.2% Tween-20) and incubated with secondary antibody for 1 hour at room temperature. Slides were mounted using Vectashield with DAPI (Vector Labs). Slides were imaged with a Nikon Eclipse 90i microscope with Volocity software (Perkin Elmer).

### Antibodies

For ChIP-seq the following antibodies were used: H3K27me3 (GeneTex GTX60892), RAD21 (Abcam ab992). For Western blot: RAD21 (Millipore 05-908), WAPL (Abcam ab70741), GAPDH (CST 5174T). For immunofluorescence: HA (Roche 11867423001).

### ChIP-seq and calibrated ChIP-seq

ChIP-seq was performed as described previously with minor modifications (Del Rosario et al., 2019). Briefly, cells were trypsinized, resuspended in 1 million cells/mL of media and crosslinked with 1% formaldehyde for 10 min at room temperature with rotation. Cells were then quenched with 0.2 M glycine for 5 min. Crosslinked cells were washed once with cold PBS, pelleted, and snap frozen in an EtOH/dry ice bath. Cell pellets were stored at -80C until further processing. 5-10 million cross-linked cells per IP were thawed on ice and resuspended in 300ul Lysis Buffer 1 (25mM HEPES pH 7.2-7.5, 25mM EDTA, 0.5% SDS). Nuclei were sonicated (Qsonica Q800 Sonicator) in polystyrene tubes at 40% power setting, 30s ON/30s OFF for a total sonication (ON) time of 6 min. After sonication, 1.2 mL of Lysis Buffer 2 (25mM HEPES pH 7.2-7.5, 187.5mM NaCl, 1.25% Triton X-100, 0.625% Na-deoxycholate, 6.25% glycerol, 1x cOmplete EDTA-free protease inhibitor cocktail) was added. 30ul of Dynabeads Protein G (Thermo Fisher Scientific) per IP were washed 3 times with ChIP Wash Buffer 1 (25mM HEPES pH 7.2-7.5, 150mM NaCl, 5mM EDTA, 1% Triton X-100, 0.5% Na-deoxycholate, 5% glycerol, 1x cOmplete EDTA-free protease inhibitor cocktail) and resuspended in 60ul ChIP Wash Buffer 1. Each IP was blocked with 20ul washed beads for 2 hours at 4C with rotation. The other 40ul of washed beads were incubated with antibody (5ul of RAD21 antibody or 2ul of H3K27me3 antibody/IP) for 2 hours at 4C. 5% input was taken from blocked lysates. Beads conjugated to antibody were added to blocked lysates and IP allowed to proceed overnight at 4C with rotation. IPs were then washed 3 times with cold ChIP Wash Buffer 1, 3 times with cold ChIP Wash Buffer 2 (25mM HEPES pH 7.2-7.5, 300mM NaCl, 5mM EDTA, 1% Triton X-100, 0.5% Na-deoxycholate, 5% glycerol, 1x cOmplete EDTA-free protease inhibitor cocktail). And 1 time with cold TENG (50mM Tris-HCl pH 8, 1mM EDTA, 50mM NaCl, 5% glycerol, 1x cOmplete EDTA-free protease inhibitor cocktail). All washes were performed by inverting tubes and setting aside for ∼2 minutes per wash. IPs were then resuspended in 500ul TENG with 100ug RNAse A (Thermo Fisher Scientific) and incubated for 30 min at 37C. IPs were then washed 2 times with cold ChIP Wash Buffer 2 and 2 times with cold TENG without protease inhibitors. IPs were resuspended in 200ul elution buffer (50mM Tris-HCl pH 7.5, 10mM EDTA, 1% SDS) and incubated 15 min at 65C. Eluate was then transferred to a new tube and 198ul TE buffer (Ambion) and 40ug Proteinase K (Sigma) added. Inputs were thawed, TE buffer added to equalize volume to IPs and 200ul elution buffer and 40ug Proteinase K added. Inputs and IPs were then incubated for 2h at 45C and crosslinks reversed overnight at 65C. DNA was purified with phenol-chloroform extraction and resuspended in 50ul TE. Efficiency of ChIP was assayed via qPCR, with samples displaying over 10-fold enrichment of positive regions over negative control prepared for sequencing. Libraries were prepared from ∼500ng of input or IP DNA using the NEBNext® Ultra™ II DNA Library Prep Kit for Illumina® (NEB) according to kit instructions. Experiments were performed in two biological replicates. Input and ChIP-seq libraries were sequenced on Illumina HiSeq, generating 20-50 million paired end 50 or 150 nucleotide reads per sample.

Calibrated ChIP-seq was performed using the same protocol as ChIP-seq except that 50,000 HEK293 cells per 5 million mouse cells were mixed in at the initial lysis step. Cells were counted and equal numbers of cells used per IP.

### ChIP-seq and calibrated ChIP-seq analysis

For uncalibrated ChIP-seq, input and ChIP-seq reads were aligned to the 129S1/SvJm (mus) and CAST/Eih (cas) genome as previously described (Kung et al., 2015; Minajigi et al., 2015). PCR duplicates were removed and uniquely mapped reads used for further analysis. Fpm-normalized bigwig tracks were generated for all uniquely mapped (comp), mus-specific (mus), and cas-specific (cas) reads.

Heatmaps and metaplots were generated using deepTools functions computeMatrix and plotHeatmap with the reference point as the summit of the called ChIP peaks.

Calibrated ChIP-seq analysis was performed as previously described (Hu et al., 2015). Briefly, paired-end reads were aligned to a concatenated mouse and human genome (mm9 + hg19) using Bowtie2 with “--no-mixed” and “--no-discordant” options. PCR duplicates were filtered out with SAMBAMBA (Tarasov et al., 2015). Uniquely aligned mouse reads were normalized to human spike-in by subsampling mm9 reads so that total number of hg19 reads was matched between samples and bigWig tracks generated from normalized samples. For calibration of RAD21 cChIP between RAD21-FKBP^degron^ clones, only reads within hg19 peaks (hg19 tracks combined) were counted for normalization purposes. Subsampling factors were further adjusted based on ratio of hg19 to mm9 reads in matched input samples. After normalization factor was determined, paired end reads were aligned to mus and cas genomes as previously described, except using Bowtie2 with the same parameters as above (with “--no-mixed” and “--no-discordant”) instead of Novoalign. PCR duplicates were filtered out with SAMBAMBA and allelic reads normalized using the same normalization factor as above.

Input subtracted H3K27me3 calibrated ChIP coverage profiles were generated with SPP (Kharchenko et al., 2008) with smoothing using 1kb windows recorded every 500 bp, as previously described (Simon et al., 2013). For input subtraction, inputs from DMSO and dTAG-13 treated samples were combined to ensure equal profiles were subtracted from each calibrated IP.

Heatmaps and metaplots were generated using deepTools functions computeMatrix and plotHeatmap with the reference point as the summit of the called ChIP peaks. Plots of calibrated H3K27me3 signal over upregulated or downregulated genes were calculated using “scale-regions” with the entire gene scaled to 3kb.

To compare the allelic densities of H3K27me3 over different classes of X-linked genes, we computed average densities over gene bodies using multiBigwigSummary from deepTools with default parameters. Non-expressed genes were defined as X-linked genes have 0 RPKM in at least one RNA-seq replicate of DMSO control. Genes subject to XCI were identified as the set of genes have allelic information (see RNA-seq analysis above), with escapees, Xist and Tsix excluded. Escapees were as defined previously (Wang et al., 2018). To compare allelic densities of H3K27me3 over S1 and S2 compartments, we defined S1 compartments as regions having positive value in the first Principal Component calculated in the WAPL-FKBP^degron^ DMSO control days 4-7 sample and S2 compartments as regions have negative value. S1 and S2 compartments were then binned into 100kb windows using bedtools makewindows, excluding windows that overlapped with unmappable regions as defined previously (Wang et al., 2018). We then we computed average densities over these 100kb bins using multiBigwigSummary from deepTools with default parameters.

### RNA-seq

Total RNA was extracted using TRIzol Reagent (Thermo Fisher Scientific) and mRNA isolated using the NEBNext Poly(A) Magnetic Isolation Module (NEB) according to the kit instructions. Two biological replicates of RNA-seq libraries were prepared using the NEBNext Ultra Directional RNA Library Prep Kit for Illumina (NEB) according the kit instructions. Libraries were sequenced on the Illumina HiSeq2500 (rapid run), generating 30-50 million 50 nt paired end reads per sample or the Illumina HiSeq4000, generating 16-27 million 150 nt paired end reads per sample.

### RNA-seq analysis

RNA-seq reads were aligned allele-specifically to the 129S1/SvJm (mus) and CAST/Eih (cas) genomes using TopHat2 (Kim et al., 2013) as previously described (Kung et al., 2015; Minajigi et al., 2015). PCR duplicates were removed and unique reads mapping to exons of each gene quantified by Homer (Heinz et al., 2010). DESeq 2 was used to call differentially expressed genes with a cut off of adjusted p-value < 0.1. For analyses on all genes, no FPKM threshold was applied, but genes were required to have at least 1 count in one sample. An expression threshold of FKPM > 0.5 was applied for analyses on expressed genes. Expressed genes with a minimum of 13 allelic-specific reads in at least one sample were counted as genes with allelic information (Colognori et al., 2019). For a given gene, fraction mus was calculated as the fraction of allelic reads that were mus-specific (mus / (mus+cas) reads).

Log2FoldChange and log2baseMean from DESeq2 were plotted for indicated classes of genes. H3K27me3 genes were defined as genes overlapping H3K27me3 cChIP-seq peaks in their respective samples (DMSO and dTAG-13 peaks combined). H3K27me3 loop genes were defined as H3K27me3 genes overlapping H3K27me3-bound loop anchors called in RAD21-FKBP^degron^ dTAG-13 Hi-C maps (i.e., cohesin-independent H3K27me3 anchored loops). For overlap with RAD21 loop anchors, loops were called in respective DMSO Hi-C maps.

### PRO-seq library construction

PRO-Seq library construction was performed by the Nascent Transcriptomics Core at Harvard Medical School, Boston, MA. Aliquots of frozen (-80°C) permeabilized cells were thawed on ice and pipetted gently to fully resuspend. An aliquot was removed and diluted 1:10 in 1xPBS for cell counting using an automatic counter (Biorad TC-20) with counts spot checked by manual recount using a hemocytometer as needed. An aliquot of each dilution was also stained with Trypan Blue and counting repeated to determine permeabilization efficiency. For each sample, 1 million permeabilized cells were used for nuclear run-on. For normalization, 50,000 permeabilized *Drosophila* S2 cells were added to each sample of 1 million cells. Nuclear run on assays and library preparation were then performed essentially as described [Elrod, N.D., et al. (2019) *Molecular Cell* **76**: 738-752.e7] with modifications noted below.

The nuclear run-on buffer was prepared as 4X stock (20mM Tris (pH 8), 20mM MgCl2, 2mM DTT, 600mM KCl, 40uM/ea biotin-11-NTPs (Perkin Elmer), 20U SuperaseIN (Thermo)). The 4X stock was mixed 1:1 with 2% sarkosyl (Sigma) to yield 2X complete run-on mix. The 3’ adapter (RNA oligo: 5’P-GAUCGUCGGACUGUAGAACUCUGAAC-3’InvdT) was pre-adenylated prior to use (5’ DNA adenylation kit, NEB, according to the manufacturer’s instructions). Adenylated oligo was purified by ethanol precipitation, resuspended in water and quality was modification confirmed by electrophoresis on a 15% TBE-Urea gel (Novex) next to appropriate controls. The adapter concentration was adjusted to 10uM and 1uL was used for each ligation which used T4 RNA ligase 2, truncated KQ (NEB) with 15% PEG-8000 in the reaction mix. Reactions were incubated at 16°C overnight. Each ligation reaction was then diluted with 180uL of betaine binding buffer (1.42g of betaine brought to 10mL with binding buffer and sterile filtered) containing 1uL of 100uM blocking oligo (TCCGACGATCCCACGTTCCCGTGG/3InvdT/). The blocking oligo is complementary to and overhangs the 3’ end of the 3’ adapter and reduces 5’/3’ adapter dimer formation during the 5’ adapter ligation step. The blocking oligo has modifications at its 3’ end, to prevent extension by reverse transcriptase. Blocking oligo was also included in final wash solution prior to the 5’ adapter ligation and in the ligation reaction itself, both at 3-fold molar excess over the initial amount of 3’ adapter (i.e. 1uL of 30uM per reaction). The 5’ adapter ligation was performed as in Elrod except PEG-8000 was increased to 15%.

After reverse transcription, cDNA was immediately amplified for 5-cycles (“preCR” step). A PCR cocktail consisting of NEBNext Ultra II Q5 master mix (NEB) and Illumina TruSeq PCR primers (RP-1, common; and RPI-X, indexing) was added to each cDNA and amplification was done according to manufacturer’s suggested 2-step cycling conditions for NGS applications. To determine optimal library amplification, the preCRs were thawed on ice and an aliquot serially diluted. Each dilution was mixed with a PCR cocktail (Q5 DNA polymerase with optional high GC enhancer) containing 10uM/ea of Illumina universal P5 and P7 primers and amplified 15-cycles. The reactions were electrophoresed on a 2.2% agarose gel (SeaKem) cast with SYBR gold (Thermo) and analyzed to determine the optimal number of cycles for final amplification. preCRs were transferred from ice to a pre-heated thermal cycler block and amplified for the appropriate number of additional cycles to reach the total determined in the test amplification.

Pooled libraries were sequenced using Illumina NovaSeq S1 (PE 50) platform.

### PRO-seq analysis

Paired-end reads were aligned to a concatenated mouse and fly genome (mm9 + dm6) using Bowtie2 with “--no-mixed” and “--no-discordant” options. DMSO and dTAG-13 samples for each clone were normalized to drosophila spike-in by subsampling mm9 reads so that total number of dm6 reads was matched between samples. For RAD21-FKBP^degron^ clone G9, downsampling factors corresponded to library size (i.e., were equivalent to downsampling based on total number of reads). For RAD21-FKBP^degron^ clone C4, downsampling factors were 1 for DMSO and 0.73 for dTAG-13 although library sizes were similar, indicating that there were overall lower levels of nascent transcription in the dTAG-13 sample. Duplicates were not removed to preserve reads corresponding to TSS and promoter proximal PolII positions. Reads mapping to genes (full gene bodies) were quantified by Homer (Heinz et al., 2010), removing miRNA, snoRNA and scaRNAs. DESeq 2 was used to call differentially expressed genes with a cut off of adjusted p-value < 0.1. For analyses on all genes, no FPKM threshold was applied, but genes were required to have at least 1 count in one sample. An expression threshold of FKPM > 0.5 was applied for analyses on expressed genes.

Log2FoldChange and log2baseMean from DESeq2 were plotted for indicated classes of genes. H3K27me3 genes were defined as genes overlapping H3K27me3 cChIP-seq peaks in their respective samples (DMSO and dTAG-13 peaks combined). H3K27me3 loop genes were defined as H3K27me3 genes overlapping H3K27me3-bound loop anchors called in RAD21-FKBP^degron^ dTAG-13 Hi-C maps (i.e., cohesin-independent H3K27me3 anchored loops). For overlap with RAD21 loop anchors, loops were called in respective DMSO Hi-C maps.

For allelic analysis, PRO-seq reads not mapping to the fly genome (dm6) were aligned to mus and cas genomes using Bowtie2 with the same parameters as above (“--no-mixed” and “--no-discordant”) and downsampled using the same factors as calculated above. Expressed genes with a minimum of 13 allelic-specific reads in at least one sample were counted as genes with allelic information (Colognori et al., 2019). For CDPs, normalized mus and cas counts divided by gene length were plotted.

### *In situ* Hi-C and visualization

*In situ* Hi-C was done as described previously (Rao et al., 2014) using 2 clones for each degron line and 5 million cells per sample. See Supplemental Table 1 for Hi-C statistics.

Reads were trimmed using cutadapt with the options --adapter=GATCGATC (MboI ligation junction) and --minimum-length=20. Reads of each pair were individually mapped to the mus (Xi) and cas (Xa) reference genomes using novoalign and merged into Hi-C summary files and filtered using HOMER as previously described (Minajigi et al., 2015). Hi-C contact maps were generated from reads in the filtered HOMER mus and cas tag directories using the ‘pre’ command of Juicer tools with the genomeID mm9 (Durand et al., 2016b). The resulting Hi-C contact maps in .hic format were visualized and normalized with the ‘Coverage Square Root’ option in Juicebox version 1.8.8 (Durand et al., 2016a).

### *In situ* Hi-C analysis

Principal component analysis (PCA) was done using the runHiCpca.pl command of HOMER with the filtered HOMER Hi-C tag directory as input and the options -res 100000 -superRes 100000 - genome mm9. Pearson correlation maps were also computed by HOMER using the analyzeHiC command with the filtered HOMER Hi-C tag directory as input and the options -chr chrX -res 200000 –corr.

Loops were called from Hi-C interaction maps using the GPU version of HiCCUPS (Durand et al., 2016) with the following parameters: ‘java -jar juicer_tools_1.13.02.jar hiccups -m 512 -r 25000,50000 -k KR -f .1,.1 -p 1,1 -i 3,3 -t 0.02,1.5,1.75,2 -d 50000,100000’, adding the -- ignore_sparsity option when needed. Loops were called from merged Hi-C interaction maps for each condition (DMSO and dTAG-13). Loops were additionally filtered to require at least 27 contacts in the peak center, 2.25 enrichment over the donut neighborhood and an FDR < 0.05 in the donut neighborhood. APA was performed on loops called by HiCCUPS using Juicer (Durand et al., 2016) at 5kb resolution.

For inter-chromosomal Hox interactions, APA was performed at 100kb resolution with the -e option using an input loop list of inter-chromosomal interactions anchored at 500kb intervals centered at Hox clusters.

For super-enhancer interactions, APA was performed at 100kb resolution using an input loop list of interactions anchored at 500kb intervals centered at super-enhancers (Whyte et al., 2013). Super-enhancer intervals were split into 5 quantiles based on distance between anchors.

TAD scores were calculated as described previously. Specifically, 100-kb resolution ‘Balanced’ normalized chromosome Hi-C interaction matrices were extracted from Juicebox using the ‘dump’ function of Juicer tools (juicer dump observed KR name.hic X X BP 100000 name.KR.100kb.dump). Custom shell and R scripts were used to convert this output matrix into the cworld matrix format. Insulation scores were then calculated for the X chromosome using the resulting cworld matrix and the matrix2insulation.pl cworld script with parameters –is 500000 –ids 400000 –im iqrMean–nt 0 –ss 200000 –yb 1.5 –nt 0 –bmoe 0. As described previously, regions of insulation score minima represent regions of locally high insulation and are considered potential TAD boundaries (Crane et al., 2015). TAD scores were then calculated using insulation scores from given samples and TAD borders called previously on the Xa (Colognori et al., 2019) with the cworld script insulation2tads.pl with default parameters.

### CHART-seq

Xist CHART-seq was performed as previously described (Wang et al., 2018). Briefly, cells were trypsinized, resuspended in 1 million cells/mL of media and crosslinked with 1% formaldehyde for 10 min at room temperature with rotation. Cells were then quenched with 0.2 M glycine for 5 min. Crosslinked cells were washed once with cold PBS, pelleted, and snap frozen in an EtOH/dry ice bath. Cell pellets were stored at -80C until further processing. 25 million cross-linked cells per CHART were thawed on ice and resuspended in 1 mL cold sucrose buffer. (10 mM HEPES pH 7.5, 0.3 M sucrose, 1% Triton X-100, 100 mM potassium acetate, 0.1 mM EGTA) supplemented with 0.5 mM spermidine, 0.15 mM spermine, 1 mM DTT, 1x protease inhibitor cocktail (Sigma, P8340), 10 U/mL RNase inhibitor (Roche). The cell suspension was rotated for 10 min at 4°C, after which it was diluted with additional 2 mL cold sucrose buffer. This was added to a pre-chilled 15-mL glass Wheaton dounce tissue grinder (Thermo Fisher Scientific) that had been pre-rinsed once with RNaseZAP RNase decontamination solution (Thermo Fisher Scientific), three times with DEPC-treated water, and once with 2 mL sucrose buffer. Release of nuclei was accomplished by douncing cells 20 times with a tight pestle, and verified by staining with Trypan Blue Solution (Thermo Fisher Scientific). The nuclear suspension was then layered atop a cushion of 7.5 mL cold glycerol buffer (10 mM HEPES pH 7.5, 25% glycerol, 1 mM EDTA, 0.1 mM EGTA, 100 mM potassium acetate) supplemented with 0.5 mM spermidine, 0.15 mM spermine, 1 mM DTT, 1x cOmplete EDTA-free protease inhibitor cocktail (Roche), and 5 U/mL RNase inhibitor, and centrifuged at 1500 g for 10 min at 4°C. The purified nuclear pellet was resuspended in 6 mL PBS and further cross-linked with 3% formaldehyde for 30 min at room temp with rotation. Afterward, nuclei were pelleted by centrifugation at 1,000 g for 5 min at 4°C and washed three times with cold PBS. Nuclei were resuspended in in 1 mL cold nuclear extraction buffer (50 mM HEPES pH 7.5, 250 mM NaCl, 0.1 mM EGTA, 0.5% sarkosyl, 0.1% sodium deoxycholate, 5 mM DTT, 10 U/mL RNase inhibitor) and incubated for 10 min at 4°C with rotation. Nuclei were then centrifuged at 400 g for 5 min at 4°C and resuspended in 230 μL cold sonication buffer (50 mM HEPES pH 7.5, 75 mM NaCl, 0.1 mM EGTA, 0.5% sarkosyl, 0.1% sodium deoxycholate, 0.1% SDS, 5 mM DTT, 10 U/mL RNase inhibitor) to a final volume of ∼270 μL. Nuclei were sonicated for 5 min at 4°C using a Covaris E220e (140W peak incident power, 10% duty factor, 200 cycles/burst). Sonicated chromatin was centrifuged at 16,000 g for 20 min at 4°C to remove debris. The supernatant (∼220 μL) was mixed with 110 μL sonication buffer to a final volume of ∼330 μL, which was split into two CHART reactions (Xist capture using antisense probes and control capture using sense probes).

For two CHART reactions, 600 μL MyOne Streptavidin C1 beads (Thermo Fisher Scientific) were used, which were pre-washed with 1 mL DEPC-treated water, blocked with 600 μL blocking buffer (33% sonication buffer, 67% 2x hybridization buffer, 500 ng/μL yeast RNA [Thermo Fisher Scientific], 1% BSA) at room temp for 1 hr, washed with 1 mL DEPC-treated water again, and resuspended in 610 μL 1x hybridization buffer (33% sonication buffer, 67% 2x hybridization buffer [see below for composition]). For each CHART, 160 μL chromatin was mixed with 320 μL 2x hybridization buffer (50 mM Tris pH 7.0, 750 mM NaCl, 1% SDS, 1 mM EDTA, 15% formamide, 1 mM DTT, 1 mM PMSF, 1x cOmplete EDTA-free protease inhibitor cocktail, 100 U/mL RNase inhibitor) and pre-cleared with 60 μL pre-blocked MyOne Streptavidin C1 beads at room temp for 1 hr with rotation. Pre-cleared chromatin was separated from beads, with 1% saved as “input” sample. The remaining pre-cleared chromatin (∼500 μL) was mixed with 36 pmol of either antisense (Xist-targeting) or sense (control) biotinylated capture probes. We used a pool of 9 capture probes, which were selected from the 11 probes described previously (Simon et al., 2013), excluding probes X.3 and X.8. Hybridization was carried out at 37°C for 4 hr followed by incubation with 240 μL blocked MyOne Streptavidin C1 beads at 37°C for 1 hr. The beads were washed once with 1x hybridization buffer (33% sonication buffer, 67% 2x hybridization buffer) at 37°C for 10 min, five times with wash buffer (10 mM HEPES pH 7.5, 150 mM NaCl, 2% SDS, 2 mM EDTA, 2 mM EGTA, 1 mM DTT) at 37°C for 5 min, and twice with elution buffer (10 mM HEPES pH 7.5, 150 mM NaCl, 0.5% NP-40, 3 mM MgCl2, 10 mM DTT) at 37°C for 5 min. 10% of the final wash was saved as an “RNA capture” sample to estimate the efficiency of Xist RNA capture afterward by qRT-PCR. CHART-enriched DNA was eluted twice in 200 μL elution buffer supplemented with 5 U/μL RNase H (New England BioLabs) at 37°C for 30 min. An “input” sample and CHART DNA were treated with 0.5 mg/mL RNase A (Thermo Fisher Scientific) at 37°C for 1 hr and then with 1% SDS, 10 mM EDTA, and 0.5 mg/mL proteinase K (Sigma) at 55°C for 1 hr. Reversal of crosslinks was performed by addition of 150 mM NaCl (final 300 mM) and incubation at 65°C overnight. DNA was purified by phenol-chloroform extraction and further sheared to < 500-bp fragments using Covaris E220e (140W peak incident power, 10% duty factor, 200 cycles/burst). Sonicated DNA was purified using 1.8x Agencourt AMPure XP beads (Beckman Coulter). Input and CHART-seq libraries were prepared using the NEBNext® Ultra™ II DNA Library Prep Kit for Illumina® (NEB) according to kit instructions. Libraries were sequenced on Illumina, generating ∼50 million 150-nt paired-end reads per sample.

### CHART-seq analysis

Input and ChIP-seq reads were aligned to the 129S1/SvJm (mus) and CAST/Eih (cas) genome as previously described (Kung et al., 2015). PCR duplicates were removed and uniquely mapped reads used for further analysis. Fpm-normalized bigwig tracks were generated for all uniquely mapped (comp), mus-specific (mus), and cas-specific (cas) reads. CHART libraries were sampled to equal read depth and input subtracted CHART coverage profiles were generated with SPP (Kharchenko et al., 2008) with smoothing using 1kb windows recorded every 500 bp, as previously described (Simon et al., 2013). After scaling, we computed base 2 logarithm of the ratio of the coverage of DMSO control divided by that of dTAG-13 using bigWigCompare from deepTools with --binSize 1000 and default parameters otherwise. To compare Xist densities over centromeric, middle, and telomeric regions, we split the first 30Mb, middle 100Mb and last 30Mb of the X chromosome into 100kb windows using bedtools makewindows, excluding windows that overlapped with unmappable regions as defined previously (Wang et al., 2018) and computed average Xist densities over these bins using multiBigwigSummary from deepTools with default parameters. To compare the densities of Xist over different classes of X-linked genes, we computed average densities over gene bodies using multiBigwigSummary from deepTools with default parameters. Non-expressed genes were defined as X-linked genes have 0 FPKM in at least one RNA-seq replicate of DMSO control. Genes subject to XCI were identified as the set of genes have allelic information (see RNA-seq analysis above), with escapees, Xist and Tsix excluded. Escapees were as defined previously (Wang et al., 2018). To compare densities of Xist over S1 and S2 compartments, we defined S1 compartments as regions having positive value in the first Principal Component calculated in the WAPL-FKBP^degron^ DMSO control days 4-7 sample and S2 compartments as regions have negative value. S1 and S2 compartments were then binned into 100kb windows using bedtools makewindows, excluding windows that overlapped with unmappable regions as defined previously (Wang et al., 2018). We then we computed average densities over these 100kb bins using multiBigwigSummary from deepTools with default parameters.

## SUPPLEMENTAL NOTE

### Long-term WAPL or CTCF degradation in ES cells induces expression of “2C-like” genes

We wondered whether the same Polycomb target genes were dysregulated in the WAPL and RAD21 degron cells. Although one might expect the DEGs to overlap but be dysregulated in opposite direction (e.g., down in RAD21-FKBP^degron^ cells, up in WAPL-FKBP^degron^ cells), there was little overlap between the two sets of dysregulated genes. In our day 7 datasets, 0 genes were DE in opposite directions in the RAD21 and WAPL degrons, while 8 genes were upregulated and 13 genes downregulated in both. On the other hand, this may not be surprising due to our WAPL degradation time course (3 days) being much longer than that of RAD21 (8 hours). We therefore wondered whether WAPL degradation would better phenocopy long term degradation of other architectural factors. Accordingly, we examined a published CTCF auxin-AID degron RNA-seq dataset, in which CTCF had been degraded for 4 days in mESCs (Nora et al., 2017). Interestingly, there was high overlap between genes DE in the same direction after WAPL or CTCF degradation (959 upregulated in both, 467 downregulated in both), with less of an overlap between genes DE in opposite directions (301 upregulated after CTCF degradation and downregulated after WAPL degradation, 74 downregulated after CTCF degradation and upregulated after WAPL degradation).

Among the genes strongly upregulated upon both WAPL and CTCF degradation were the *Zscan4* and *Usp17l* families (**Figure S3E**), markers of the two-cell embryonic development stage (Falco et al., 2007; Macfarlan et al., 2012). Subpopulations of mESCs in cell culture have been found to express such a “2C-like” gene signature, providing an explanation for why these genes could be expressed in the ES cell degron lines of us and others (Kolodziejczyk et al., 2015; Macfarlan et al., 2012). Notably, *Usp17l* is a family of de-ubiquitinating enzymes (Reyes-Turcu et al., 2009). As both dTAG and the auxin-AID degron system rely on tethering degron-tagged proteins to ubiquitin ligases, upregulation or selection of a subpopulation expressing these “2C- like” genes could reflect use of degrons in mESCs for extended time periods. Alternatively, a recent study implicated cohesin in regulation of 2-cell genes (Zhang et al., 2020), supporting that the upregulation we observe in the case of CTCF and WAPL degradation could be a consequence of toggling cohesin localization on chromatin. We emphasize that, although “2C-like” genes were strongly upregulated, their overall expression level remained relatively low and WAPL-FKBP^degron^ ES cells were still able to differentiate and efficiently degrade WAPL upon addition of dTAG-13 (Figure 1B). Therefore, the percent of “2C-like” cells in the population likely remained negligible.

